# Preferential cholinergic excitation of corticopontine neurons

**DOI:** 10.1101/182667

**Authors:** Arielle L. Baker, Ryan J. O’Toole, Allan T. Gulledge

## Abstract

Pyramidal neurons in layer 5 of the neocortex comprise two broad classes of projection neurons: corticofugal neurons, including corticopontine (CPn) neurons, and intratelencephalic neurons, including commissural/callosal (COM) neurons. These non-overlapping neuron subpopulations represent discrete cortical output channels contributing to perception, decision making, and behavior. CPn and COM neurons have distinct morphological and physiological characteristics, and divergent responses to modulatory transmitters such as serotonin and acetylcholine (ACh). To better understand how ACh regulates cortical output, in slices of mouse prefrontal cortex (PFC) we compared the responsivity of CPn and COM neurons to transient exposure to exogenous or endogenous ACh. In both neuron subtypes, exogenous ACh generated qualitatively similar biphasic responses in which brief hyperpolarization was followed by longer-lasting enhancement of excitability. However, cholinergic inhibition was more pronounced in COM neurons, while excitatory responses were larger and longer lasting in CPn neurons. Similarly, optically triggered release of endogenous ACh from cholinergic terminals preferentially and persistently (for ~40 s) enhanced the excitability of CPn neurons, but had little impact on COM neurons. Cholinergic excitation of CPn neurons involved at least three distinct ionic mechanisms: activation of a calcium-sensitive but calcium-permeable nonspecific cation conductance, suppression of K_v_7 channels (the “M-current”), and activation of the calcium-dependent nonspecific cation conductance underlying afterdepolarizations. Our results demonstrate projection-specific selectivity in cholinergic signaling in the PFC, and suggest that transient release of ACh during behavior will preferentially promote corticofugal output.

## Introduction

In the mammalian prefrontal cortex (PFC), acetylcholine (ACh) is a neurotransmitter that facilitates many cognitive functions, including attentional processes such as cue detection (Klinkenberg et al., 2011; Parikh and Sarter, 2008). For instance, loss of cholinergic input to the PFC impairs attention (Dalley et al., 2004; McGaughy et al., 2002; McGaughy et al., 1996; Newman and McGaughy, 2008), whereas enhancement of cholinergic signaling in rodents (Kolisnyk et al., 2013a), primates (Lange et al., 2015), and humans (Bentley et al., 2004) can improve performance in attention tasks. However, the specific cellular and circuit-based mechanisms by which cholinergic input to the PFC facilitates attention and other cognitive processes remains obscure.

Within the neocortex, ACh acts through a variety of metabotropic and ionotropic ACh receptors differentially expressed in subclasses of cortical neurons, including pyramidal neurons that provide the bulk of cortical output, and GABAergic interneurons that comprise local inhibitory circuits. Pyramidal neurons in layer 5 of the rodent medial PFC (mPFC) respond to ACh primarily via M1-type muscarinic ACh receptors (mAChRs; Gulledge et al., 2009) that trigger calcium release from internal calcium stores and activate SK-type calcium-activated potassium channels (Gulledge and Stuart, 2005), while simultaneously engaging voltage-dependent nonspecific cation conductances (Andrade, 1991; Haj-Dahmane and Andrade, 1996; Yan et al., 2009) that facilitate calcium influx from the extracellular space (Dasari et al., 2017).

However, cortical projection neurons are not homogenous, and in layer 5 comprise two broad, non-overlapping subpopulations that provide parallel output channels from the cortex: corticopontine (CPn) neurons projecting to the brainstem, and commissural/ callosal (COM) neurons that provide output to the contralateral cerebral hemisphere (Morishima and Kawaguchi, 2006). In addition to having distinct morphological and physiological properties (Dembrow et al., 2010; Morishima and Kawaguchi, 2006), CPn and COM neurons differ in their responses to neuromodulatory input (Dembrow and Johnston, 2014). For example, serotonin (5-HT) inhibits CPn neurons via G_i/o_-coupled 5-HT_1A_ receptors, while selectively exciting COM neurons via activation of G_q_-coupled 5-HT_2A_ receptors (Avesar and Gulledge, 2012). On the other hand, CPn neurons are preferentially excited by a2 adrenergic (Dembrow et al., 2010) and dopamine D2 (Gee et al., 2012) receptors, while COM neurons are preferentially excited via activation of dopamine D1 receptors (Seong and Carter, 2012). In terms of cholinergic modulation, layer 5 pyramidal neurons in the mPFC broadly express G_q_-coupled M1 subtype mAChRs and exhibit enhanced excitability in the continuous presence of muscarinic agonists (Gulledge et al., 2009), with responses to tonic mAChR activation being more robust in CPn neurons (Dembrow et al., 2010).

The purpose of this study was to determine whether transient cholinergic stimulation, including phasic release of endogenous ACh from cholinergic terminals, differentially influences the excitability of CPn and COM neurons in the mPFC. Our results reveal a dichotomy in transient cholinergic signaling in these two projection neuron populations that is consistent with the cell-type specificity of cholinergic responses in the mouse auditory cortex (Joshi et al., 2016). We further identify three distinct ionic mechanisms that contribute to cholinergic excitation of CPn neurons. Overall, our results suggest that phasic release of ACh in the PFC, as occurs *in vivo* during cue detection tasks (Sarter et al., 2016), will preferentially and persistently enhance corticofugal output.

## Materials and Methods

### Ethical approval

Experiments were performed using female and male 6-to 10-week-old C57BL/6J wild-type (stock #013636), ChAT-ChR2-YFP (#014546), or ChAT-IRES-Cre (#006410) x Ai32 (#012569) mice *(Mus musculus)* obtained from the Jackson Laboratory and cared for according to procedures approved by the Institutional Animal Care and Use Committee of Dartmouth College. Animals were bred in facilities accredited by the Association for Assessment and Accreditation of Laboratory Animal Care, and maintained on a 12 h/12 h light-dark cycle with food and water *ad libitum.*

### Retrograde labeling

Red or green fluorescent beads (Retrobeads, Lumafluor Inc.) were injected unilaterally into either the pons (to label ipsilaterally projecting CPn neurons) or prelimbic cortex (to label contralaterally projecting COM neurons) using age-appropriate coordinates (Paxinos and Franklin, 2004). Animals were continuously anesthetized with vaporized isoflurane (~2%) during surgeries in which a craniotomy was made and a microsyringe lowered into the brain region of interest. 200-600 nL of undiluted Retrobead solution (Lumafluor, Inc.) was injected at a rate of 0.015 µL/min. After injection, the microsyringe was held in place for ~5 min before being slowly withdrawn. Animals were allowed to recover from surgery for at least 48 (for COM injections) or 72 (for CPn injections) hours before use in electrophysiological experiments. Locations of bead injections were confirmed in coronal sections of the mPFC or brainstem.

### Slice preparation

Animals were anesthetized with vaporized isoflurane, decapitated, and brains rapidly removed into artificial cerebral spinal fluid (aCSF) composed of the following (in mM): 125 NaCl, 25 NaHCO_3_, 3 KCl, 1.25 NaH_2_PO_4_, 0.5 CaCl_2_, 6 MgCl_2_, and 25 glucose (saturated with 95% O_2_ / 5% CO_2_). Coronal brain slices (250 µm thick) of the mPFC were cut using a Leica VT 1200 slicer and stored in a holding chamber filled with aCSF containing 2 mM CaCl_2_ and 1 mM MgCl_2_. Slices were maintained in the holding chamber for 1 hour at 35 °C, and then at room temperature (~27 °C) until use in experiments.

### Electrophysiology

Slices were transferred to a recording chamber continuously perfused (~7 ml/min) with oxygenated aCSF heated to 35-36 °C. Labeled CPn or COM pyramidal neurons were visualized with epifluorescence (470 or 530 nm LEDs) via a 60x water-immersion objective. Patch pipettes (5-7 MQ) were normally filled with a solution containing the following (in mM): 135 K-gluconate, 2 NaCl, 2 MgCl_2_, 10 HEPES, 3 Na_2_ATP, and 0.3 NaGTP, pH 7.2 with KOH. In some experiments K-gluconate was replaced with BAPTA (a mixture of tetrapotassium and tetraacetic salts) to produce pipette solutions containing 10 or 30 mM BAPTA. Data were acquired using a BVC-700 amplifier (Dagan Corporation.) connected to an ITC-18 digitizer (HEKA) driven by AxoGraph software (AxoGraph Scientific). Membrane potentials were sampled at 25 kHz, filtered at 5 kHz, and corrected for the liquid junction potential of +12 mV.

### Cholinergic stimulation

To measure cholinergic responses in CPn and COM neuron excitability, we transiently delivered ACh to targeted neurons during periods of continuous action potential generation (~6-8 Hz) evoked by somatic DC current injection. For exogenous application, ACh was dissolved in aCSF (to 100 µM) and focally applied (100 ms at ~10 PSI) from a patch-pipette positioned near the soma of the recorded neuron. Release of endogenous ACh in slices from ChAT-ChR2-YFP or ChAT-IRES-Cre / Ai32 mice was triggered with wide-field flashes of blue light (470 nm, 5 ms) from an LED (Thor Labs; ~3.5 mW). Inhibitory responses to ACh were quantified as the duration of action potential cessation following the cholinergic stimulus, while excitatory responses to ACh were quantified as the peak increase in instantaneous spike frequency (ISF) relative to the average baseline ISF. Durations of excitatory responses were quantified by resampling ISF plots at 2 Hz and identifying the timing at which the ISF dropped below the mean baseline ISF.

### Pharmacological manipulations

In some experiments, apamin (100 nM), atropine (1 µM), XE991 (10 µM), cadmium (200 µM), or kyurenate (3 mM) and gabazine (10 µM), were bath applied for 5-10 minutes before ACh application or exposure to blue light. In experiments in which intracellular calcium was chelated with BAPTA, experiments started at least 10 minutes after establishment of whole-cell recording. For experiments using nominally calcium-free aCSF, CaCl_2_ was replaced with equimolar MgCl_2_. In some experiments extracellular KCl was reduced to 0.5 mM, with NaCl being raised to 127.5 mM. Most drugs were obtained from Sigma-Aldrich. XE991 was obtained from Tocris and Cayman Chemical. Brain slices exposed to drug treatment were not used for more than one experiment.

### Statistical analyses

Unless otherwise noted, data are presented as mean ± standard error of the mean (SEM), and were assessed with either a Student’s t-test (two-tailed, paired or unpaired) or a one-way ANOVA (two-tailed, repeated measures with Šidák corrected post-tests, where appropriate,) using Wizard for Mac version 1.9 (Evan Miller). Data in Tables 1 and 2 show aggregate data from baseline ACh responses in non-BAPTA-containing neurons, and presented as means ± standard deviations (SD). Significance was defined as *p* < 0.05.

**Table 1.** Physiology of layer 5 projection neurons. Table 1 compares resting membrane potential (V_M_), input resistance (R_N_), and the percent rebound sag following a current-induced hyperpolarization (% sag) in CPn and COM projection neurons from the indicated genotypes. Data are aggregated baseline data from all experiments utilizing regular (non-BAPTA-containing) intracellular solution, and presented as means ± standard deviations. No significant differences were observed between genotypes (ANOVAs), but all physiological parameters were significantly different between CPn and COM neurons within genotypes (Student’s t-tests).

| Projection subtype | Mouse genotype | n | V<sub>M</sub> (mV) | R<sub>N</sub> (MΩ) | % sag |
| --- | --- | --- | --- | --- | --- |
| CPn | Wild-type C57/BL6 | 90 | -78 ± 3 | 58 ± 29 | 17 ± 5 |
|  | ChAT-ChR2 | 70 | -78 ± 2 | 62 ± 21 | 17 ± 4 |
|  | ChAT-cre/Ai32 | 10 | -78 ± 3 | 58 ± 25 | 17 ± 4 |
| p value - within CPn (ANOVA) |  |  | 0.689 | 0.603 | 0.989 |
| COM | Wild-type C57/BL6 | 26 | -84 ± 3 | 110 ± 57 | 7 ± 4 |
|  | ChAT-ChR2 | 41 | -82 ± 5 | 131 ± 47 | 7 ± 5 |
|  | ChAT-cre/Ai32 | 9 | -83 ± 4 | 145 ± 37 | 7 ± 4 |
| p value - within COM (ANOVA) |  |  | 0.286 | 0.114 | 0.800 |
| p value - CPn vs COM (Student's t-test) |  | < 0.001 for each genotype comparison |  |  |  |

**Table 2.** Cholinergic responses in CPn and COM neurons across sex. Table 2 compares inhibitory and excitatory responses to exogenous and endogenous ACh in CPn and COM projection neurons from male and female mice from experiments in baseline conditions using regular (non-BAPTA-containing) intracellular solution. Data presented as means ± standard deviations. Within both sexes, CPn and COM were significantly different (Student’s t-tests) in all measures. No significant differences were observed between male and female mice for any parameter (Student’s t-tests), and all responses were significantly different between CPn and COM neurons in pooled data from both sexes (Student’s t-tests).

| Projection subtype | Sex | Duration of inhibition (s) | Peak excitation (% $\Delta$ Hz) | Peak excitation (% $\Delta$ Hz) |
| --- | --- | --- | --- | --- |
|  |  |  | Exogenous | Endogenous |
| CPn | Female | $0.7 \pm 0.4$<br>(n = 48) | $166 \pm 87$<br>(n = 48) | $32 \pm 15$<br>(n = 25) |
| | Male | $0.8 \pm 0.5$<br>(n = 34) | $188 \pm 70$<br>(n = 34) | $23 \pm 12$<br>(n = 13) |
| | All CPn | $0.7 \pm 0.4$<br>(n = 82) | $171 \pm 78$<br>(n = 82) | $28 \pm 15$<br>(n = 25) |
| p value - within CPn, female vs male (Student's t-test) |  | 0.61 | 0.10 | 0.06 |
| COM | Female | $1.7 \pm 0.8$<br>(n = 18) | $76 \pm 43$<br>(n = 18) | $10 \pm 7$<br>(n = 7) |
| | Male | $1.4 \pm 0.6$<br>(n = 17) | $78 \pm 32$<br>(n = 17) | $12 \pm 4$<br>(n = 4) |
| | All COM | $1.5 \pm 0.7$<br>(n = 35) | $77 \pm 37$<br>(n = 35) | $11 \pm 6$<br>(n = 11) |
| p value - within COM, female vs male (Student's t-test) |  | 0.16 | 0.97 | 0.54 |
| p value - within female, CPn vs. COM (Student's t-test) |  | < 0.001 | < 0.001 | < 0.001 |
| p value - within male, CPn vs. COM (Student's t-test) |  | 0.002 | < 0.001 | 0.023 |
| p value - All CPn vs COM (Student's t-test) |  | < 0.001 | < 0.001 | < 0.001 |

## Results

### Exogenous ACh preferentially excites CPn neurons

We initially tested the cell-type-specificity of phasic cholinergic signaling using targeted whole-cell recordings of labeled CPn and COM neurons in layer 5 of the prelimbic region of the mouse mPFC. Relative to COM neurons (n = 26), CPn neurons (n = 53) had more depolarized resting membrane potentials (-77 ± 2 vs 84 ± 3 mV for CPn and COM neurons, respectively; means ± SD; *p* < 0.001, Student’s t-test), lower input resistances (R_N_; 63 ± 35 vs 110 ± 57 MΩ, respectively; *p* < 0.001), and greater HCN-channel-mediated sag potentials (18 ± 5 vs 7 ± 4%, respectively; *p* < 0.001). Similar projection-specific physiological differences were observed in CPn and COM neurons from mice expressing ChR2 in cholinergic neurons (Table 1), but no physiological differences were observed within projection neurons across genetic models or between male and female mice (Tables 1 and 2).

To compare cholinergic responses in CPn and COM neurons, we focally applied exogenous ACh (100 µM, 100 ms) to neurons during periods of spontaneous action potential generation (~7 Hz) in response to suprathreshold DC current injection. In both neuron subtypes, ACh application generated biphasic responses in which a brief cessation of action potential firing was followed by an increase in instantaneous spike frequency (ISF; Figure 1A). However, relative to responses in COM neurons (n = 35), cholinergic inhibition was of shorter duration, and excitation of greater magnitude, in CPn neurons (n = 47; Figure 1B). Cessations of action potential generation lasted for 0.9 ± 0.1 s and 1.6 ± 0.1 s in CPn and COM neurons, respectively (CPn vs. COM, *p* < 0.001, Student’s t-test). Following these transient inhibitory responses, ISFs rose 170 ± 12% above baseline levels in CPn neurons, but only 77 ± 6% above baseline in COM neurons (CPn vs COM, *p* < 0.001). Differences in response magnitudes in CPn and COM did not result from differences in baseline firing frequency (8.1 ± 0.2 Hz vs 6.5 ± 0.2 Hz, for CPn and COM neurons, respectively, *p* < 0.001, Student’s t-test; reflecting greater spike frequency adaptation in COM neurons, see Hattox and Nelson, 2007). Baseline ISFs were not significantly correlated with the magnitude of cholinergic excitation (R^2^ = 0.067, p = 0.08 and R^2^ = 0.036, p = 0.28 for CPn and COM neurons, respectively; linear regression; Figure 1C). Responses also did not depend on sex, as equivalent projection-specific differences in cholinergic responses were observed in CPn and COM neurons from female and male mice (Table 2).

**Figure 1.**
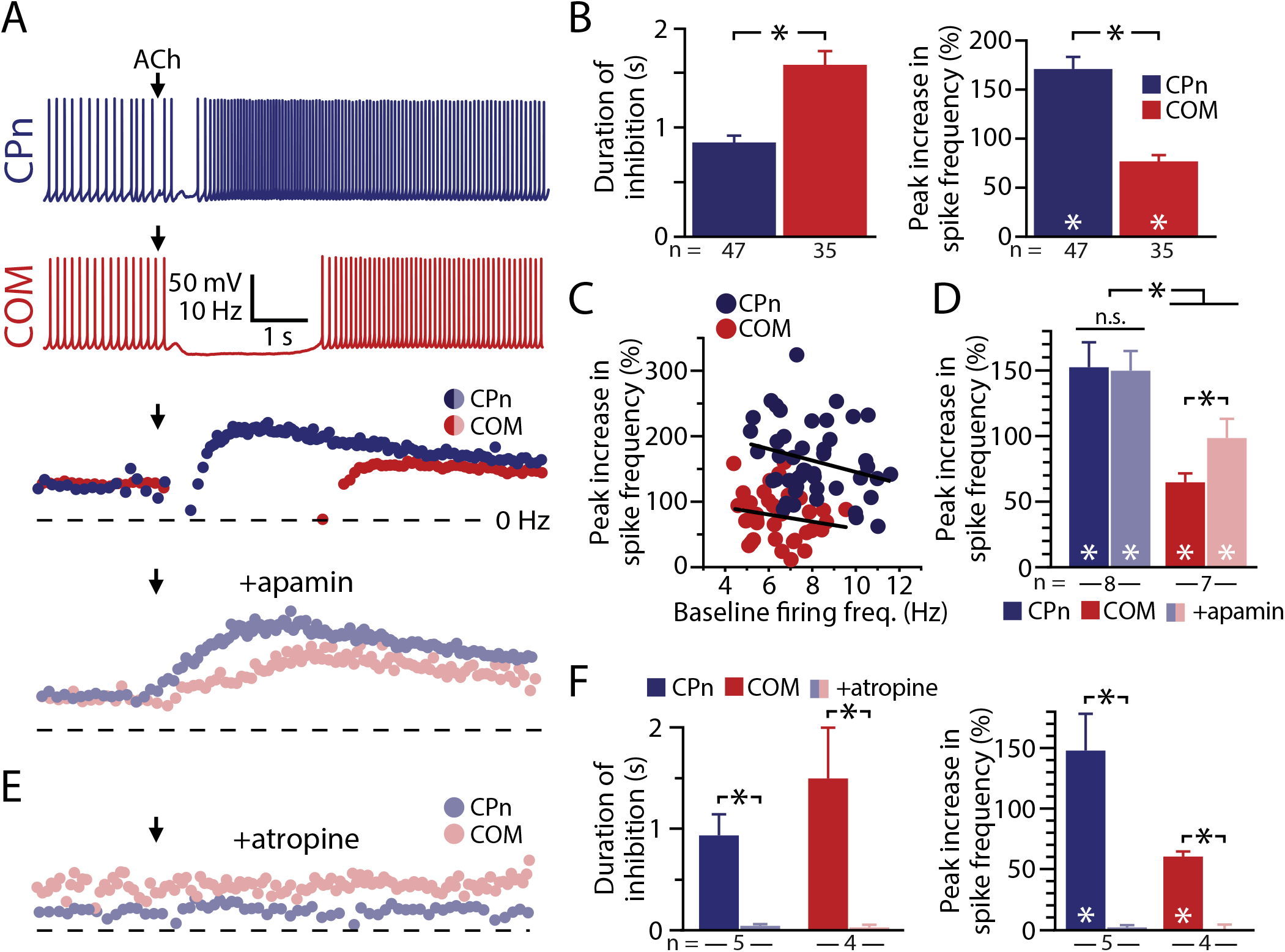
Exogenous ACh preferentially excites CPn neurons. **A)** Voltage responses of CPn (blue) or COM (red) neurons to focally applied ACh (100 µM) delivered during periods of current-induced action potential generation (top), with corresponding plots of instantaneous spike frequency (ISF) for each action potential in baseline conditions or after application of apamin (100 nM, below). Dashed line indicates 0 Hz. **B)** Comparisons of the duration of action potential inhibition (left) and the magnitude of excitatory cholinergic responses (right) in CPn (n = 47) and COM (n = 35) neurons. **C)** Plots of peak increase in spike frequency vs baseline firing frequency for CPn (blue) and COM (red) neurons. **D)** Comparison of ACh-induced peak changes in spike frequency before and after blockade of SK channels in CPn (n = 8) and COM (n = 7) neurons. **E)** Plots of ISF for CPn (blue) and COM (red) neurons in the presence of atropine (1 µM). **F)** Comparisons of inhibitory (left) and excitatory (right) responses to ACh before and after atropine treatment in CPn (n = 5) and COM (n = 4) neurons. Asterisks indicate significant differences (*p* < 0.05); white asterisks indicate significant differences from baseline firing frequencies, while black asterisks indicate significant differences between COM and CPn neurons.

To confirm that longer-lasting inhibitory responses in COM neurons were not masking and/or limiting subsequent excitation to give the appearance of preferential cholinergic excitation of CPn neurons, we measured excitatory responses to focally applied ACh before and after blocking SK channels with apamin (100 nM). Apamin eliminated cholinergic inhibitory responses in both CPn (n = 8; *p* = 0.001) and COM (n = 7; *p* = 0.014) neurons (paired Student’s t-tests), but left cholinergic excitation intact (Figure 1A). In the presence of apamin, latencies to peak cholinergic excitation decreased in COM neurons (from 3.9 ± 0.8 s to 1.9 ± 0.4 s; *p* = 0.029, paired Student’s t-test) neurons, but remained similar in CPn neurons (3.1 ± 0.3 s in baseline conditions vs 2.8 ± 0.4 s in apamin; *p* = 0.48; data not shown). Further, while blocking SK channels enhanced the magnitudes of excitatory responses in COM neurons (from 65 ± 7% to 98 ± 15% above baseline firing rates; *p* = 0.024, paired Student’s t-test), excitatory responses continued to be significantly larger in CPn neurons (150 ± 15% above baseline firing rates; *p* = 0.030 vs COM, Student’s t-test; Figure 1D). These results confirm that robust cholinergic excitation is an intrinsic property of CPn neurons independent of projection-specific differences in SK-mediated inhibition.

Both inhibitory and excitatory responses were mediated by mAChRs, as they were both sensitive to bath application of atropine (1 µM; n = 9; Figure 1E). Addition of the muscarinic antagonist eliminated all inhibitory and excitatory cholinergic responses, with durations of spike cessation being reduced from 0.9 ± 0.2 s to 0.1 ± 0 s in CPn neurons (n = 5; *p* < 0.001) and from 1.5 ± 0.5 s to 0.1 ± 0 s in COM neurons (n = 4; *p* < 0.001), while excitatory responses were reduced from 148 ± 30% to 2 ± 2% above baseline spike frequencies in CPn neurons (n = 5; *p* = 0.007) and from 60 ± 4% to 0 ± 4% above baseline firing rates in COM neurons (n = 4; *p* = 0.002; Figure 1F). These results demonstrate that transient activation of mAChRs with exogenous ACh preferentially enhances the action potential output of CPn neurons in the mouse mPFC.

### Endogenous ACh preferentially excites CPn neurons

To test the relative sensitivities of CPn and COM neurons to endogenously released ACh, we used flashes of blue (470 nm; 5 ms) light to evoke ACh release in slices of mPFC from ChAT-ChR2-YFP mice that express channelrhodopsin-2 in cholinergic neurons. We first confirmed ChR2 expression in cholinergic neurons in these animals by recording from YFP-positive cholinergic neurons in slices of the basal forebrain. Flashes of blue light reliably generated action potentials in YFP-positive neurons (Figure 2A; n = 4), while repetitive light flashes (50 Hz) generated trains of action potentials with some degree of stochastic failure that could be detected in both extracellular and whole-cell recordings (Figure 2B; n = 4).

**Figure 2.**
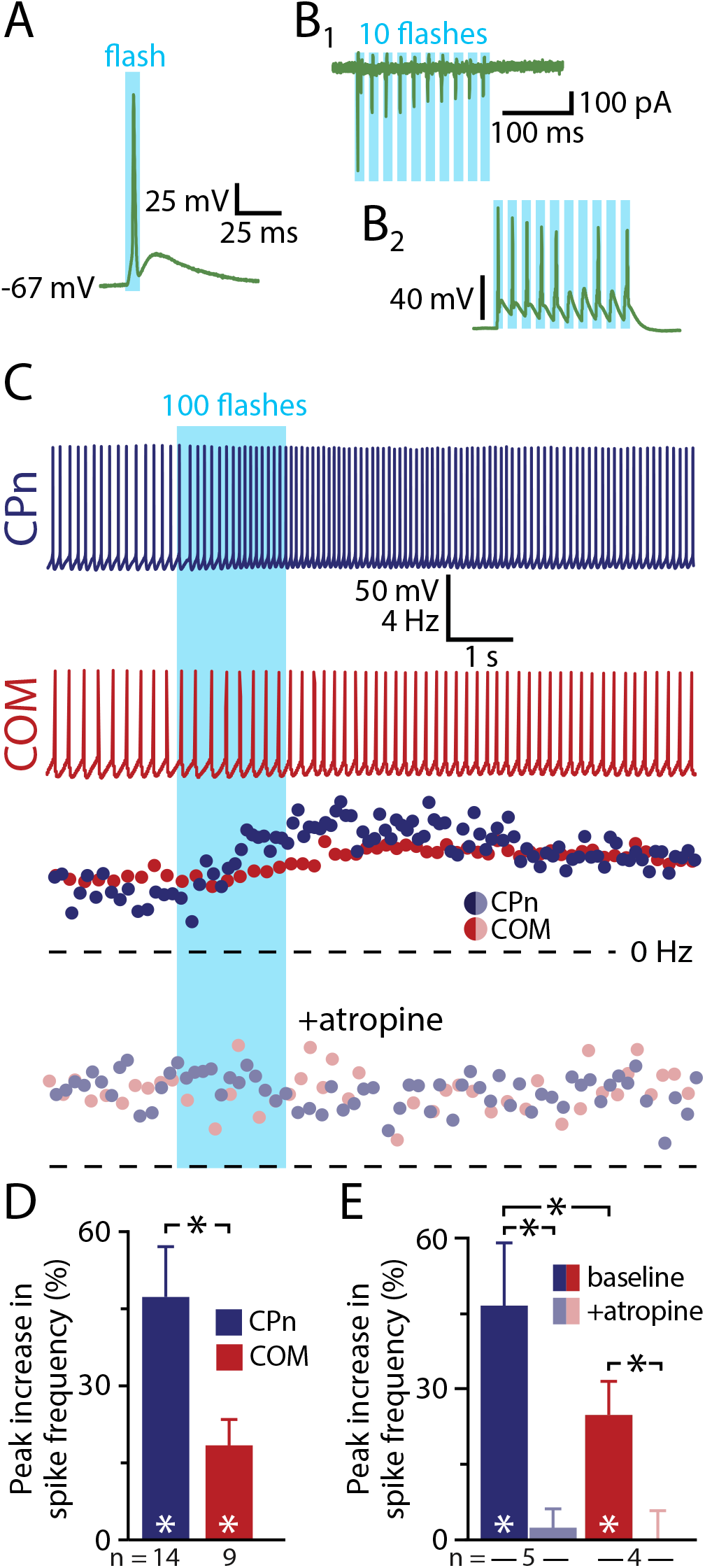
Endogenous ACh preferentially excites CPn neurons. **A, B)** Single flashes of 470 nm light (5 ms) evoked single action potentials in cholinergic neurons (n = 4) in the basal forebrain of ChAT-ChR2 mice **(A)**, while trains of light flashes (5 ms at 50 Hz) generated multiple action potentials, with some degree of failure, in extracellular recordings **(B_1_)** and in subsequent whole-cell recordings **(B_2_**; n = 4). **C)** Voltage responses and corresponding spike frequency plots for CPn (blue traces) and COM (red traces) neurons to optical activation consisting of 100 flashes of blue light (5 ms each, at 59 Hz) before (top) and after (bottom) bath application of atropine (1 µM). **D)** Comparison of peak increases in spike frequency in CPn (blue; n = 14) and COM (red; n = 9) neurons. **E)** Comparisons of cholinergic excitation before and after atropine application in a subset of CPn (n = 5) and COM (n = 4) neurons. Asterisks indicate significant differences (*p* < 0.05); white asterisks indicate significant differences from baseline firing frequencies, while black asterisks indicate significant differences between CPn and COM neurons and between experimental conditions.

To evoke endogenous ACh release in slices of mPFC, we first delivered trains of 100 flashes of blue light (5 ms each at 59 Hz) while recording from labeled CPn or COM neurons firing action potentials in response to suprathreshold DC current injection (Figure 2C). Release of endogenous ACh did not evoke significant SK responses in either projection neuron subtype, but preferentially increased the firing rates of CPn neurons by 48 ± 9% (n = 14, *p* < 0.001, paired Student’s t-test), while the firing rates of COM neurons were less affected (+20 ± 9%; n = 9, p = 0.058; Figure 2D). Overall, flash-induced increases in firing rates were larger in CPn neurons than in COM neurons (*p* = 0.007; Student’s t-test). In a subset of neurons (n = 9), we confirmed that responses to endogenous ACh were mediated by mAChRs by bath applying atropine (Figure 2C), which eliminated flash-evoked responses in both CPn (from 47 ± 13% to 2 ± 4% above baseline firing rates; n = 5; p = 0.009) and COM (from 25 ± 7% to 0 ± 8% above baseline firing rates; n = 4; p = 0.042) neurons (Figure 2E). These results confirm that release of endogenous ACh preferentially excites CPn neurons via activation of mAChRs.

We next tested whether a single flash of light, likely equating to a single ACh release event in presynaptic terminals, was sufficient to drive cholinergic responses in labeled COM and CPn neurons. To do this, we delivered periodic current steps (1.5 s duration at 0.133 Hz) that in baseline conditions generated 8.5 ± 0.1 and 8.6 ± 0.2 action potentials in COM (n = 21) and CPn (n = 18) neurons, respectively (Figure 3A). A single 5 ms flash of light, delivered at the start of the sixth current-step trial, immediately increased the number of action potentials generated in that trial. In COM neurons, the increase in action potential number was small (mean of 1.2 ± 0.3 additional spikes; p = 0.004, paired Student’s t-test) and transient, occurring only during the flash-exposed trial. On the other hand, CPn neurons exhibited larger flash-induced increases in action potential number (mean of 2.3 ± 0.3 additional spikes; *p* < 0.001) that persisted during the subsequent two trials (Figure 3B). The cholinergic origin of flash-evoked excitatory responses was confirmed in a subset of neurons exposed to atropine (1 µM; Figure 3B, inset), which eliminated single-flash-evoked changes in excitability in both CPn (n = 4) and COM (n = 6) neurons.

**Figure 3.**
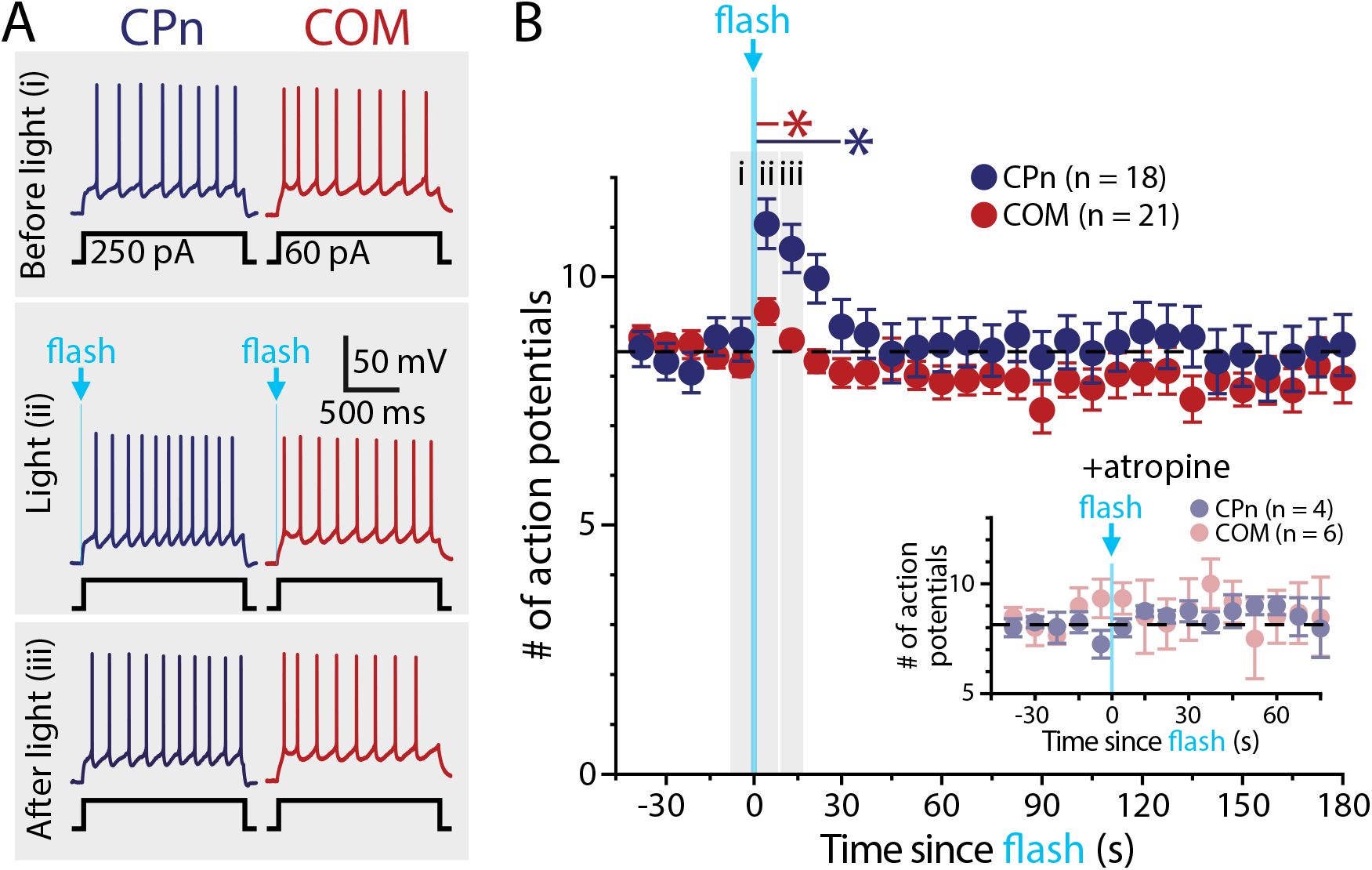
Single flash-evoked release of endogenous ACh preferentially excites CPn neurons. **A)** Periodic somatic current steps (1.5 s) generated 7-8 action potentials in baseline conditions (trial 5) in CPn and COM neurons (top). In both neuron populations, a single flash of blue light (5 ms) applied at the beginning of trial six increased the number of action potentials generated (middle), but this enhanced excitability persisted into the following trial (trial 7) only in CPn neurons (bottom). **B)** Plot of the mean number of action potentials generated by periodic current steps in populations of CPn (blue; n = 18) and COM (red; n = 21) neurons over time. Trials 5 (i), 6 (ii), and 7 (iii) seven are shaded and indicate time points of voltage traces in panel **A.** Inset: in a subset of CPn (n = 4) and COM (n = 6) neurons, the presence of atropine (1 µM) blacked flash-evoked increases in action potential generation. Asterisks indicate significant differences (*p* < 0.05); blue and red asterisks and lines indicate the duration of significant differences from baseline firing frequencies in CPn and COM neurons, respectively.

### ACh persistently excites CPn neurons

To more precisely measure the duration of persistent cholinergic excitation of layer 5 projection neurons, we repeated experiments under conditions in which neurons were continuously depolarized with suprathreshold somatic DC current injection (Figure 4A). Under these conditions, single flashes of light preferentially enhanced action potential generation in CPn neurons (mean increase in spike rate was 27 ± 3%; n = 27, *p* < 0.001, paired Student’s t-test), an effect that persisted for 23 ± 5 s after the flash (Figure 4B). Single flashes of light modestly enhanced firing rates in COM neurons by 11 ± 2% (n = 11, p < 0.001), but this effect lasted for only 7 ± 4 s. Both the magnitudes (*p* < 0.001; Student’s t-test) and durations (p = 0.048) of flash-evoked cholinergic responses were greater in CPn neurons (Figure 4C). Similarly, in experiments using focal application of exogenous ACh, CPn (n = 14) and COM (n = 10) neurons responded with brief cessation of action potential generation (0.9 ± 0.1 s vs 1.8 ± 0.4 s; p = 0.013, Student’s t-test) followed by enhanced firing frequencies (increases of 160 ± 12% and 78 ± 14%; p < 0.001) that persisted for 51 ± 7 s and 19 ± 2 s (p = 0.002) in CPn and COM neurons, respectively (Figure 4B,C). Equivalent projection-specific differences in ACh responses were observed in CPn and COM neurons from female and male mice (Table 2). These results demonstrate that transient activation of mAChRs with endogenous or exogenous ACh preferentially and persistently enhances action potential output of CPn neurons in the mouse mPFC.

**Figure 4.**
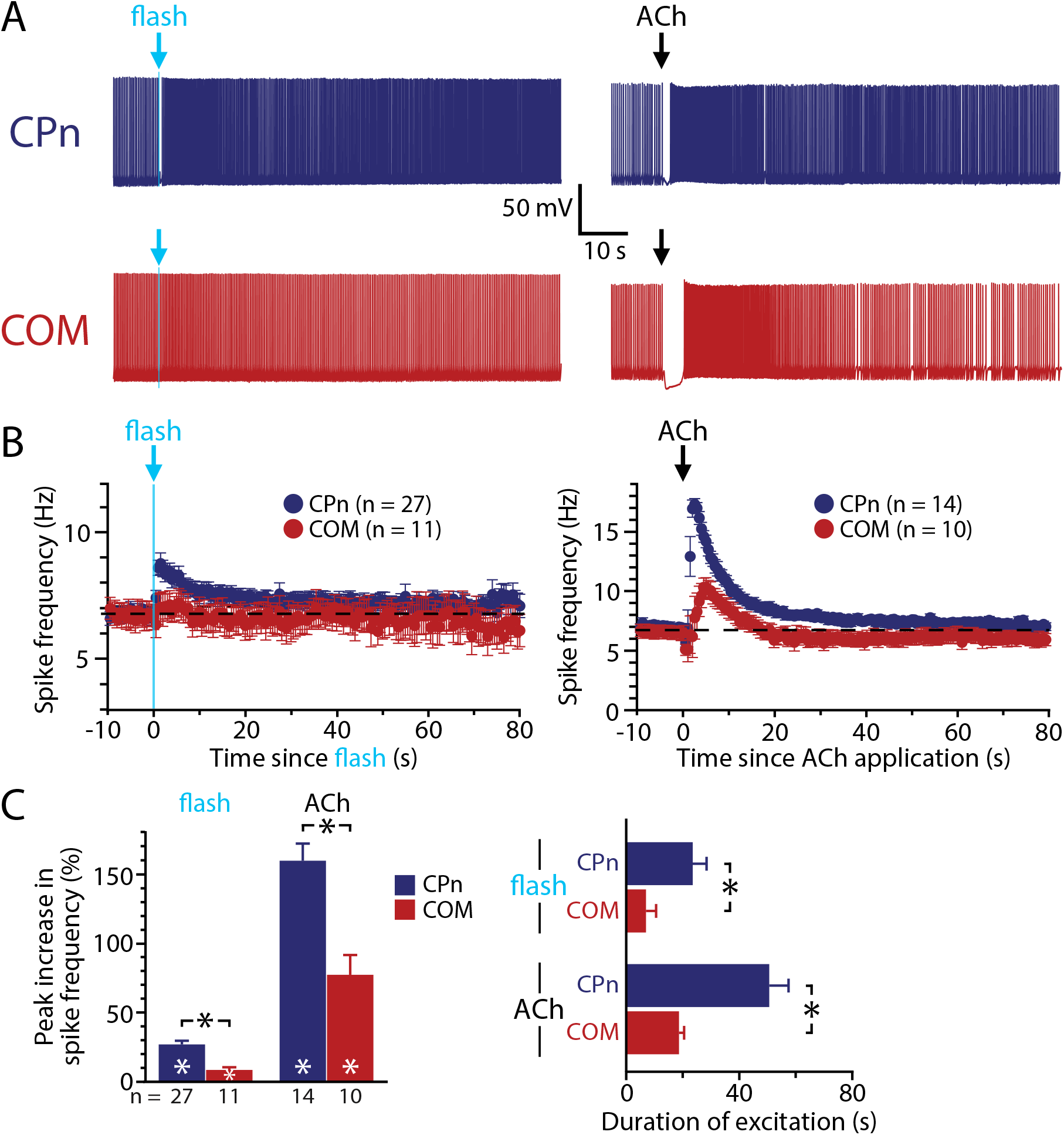
Persistence of cholinergic excitation in CPn and COM neurons. **A)** Responses to single flash-evoked release of endogenous ACh (5 ms flash, left) or exogenous ACh (100 ms, right) in CPn (blue traces; top) and COM (red traces; bottom) neurons. **B)** Plots of mean ISFs over time in populations of neurons exposed to endogenous (CPn, n = 27; COM, n = 11; left) or exogenous (CPn, n = 14; COM, n = 10; right) ACh. **C)** Comparisons of the mean increases in firing frequency following exposure to endogenous and exogenous ACh in CPn and COM neurons (left) and the duration of cholinergic excitation following endogenous or exogenous ACh exposure (right). Asterisks indicate significant differences (*p* < 0.05); white asterisks indicate significant changes from baseline firing frequencies, while black asterisks indicate significant differences between CPn and COM neurons.

### An alternative optogenetic model for ACh release

Because the ChAT-ChR2-YFP mouse line overexpresses the vesicular ACh transporter (VAChT), potentially leading to higher than normal vesicular ACh content (Kolisnyk et al., 2013a), we confirmed projection-specific cholinergic signaling in the mPFC of ChAT-Cre/Ai32 mice, an alternative animal model having ChR2 expression in cholinergic neurons, but with otherwise normal cholinergic function (Hedrick et al., 2016). As was found in tissue from ChAT-ChR2-YFP mice, flash-evoked ACh release failed to generate significant SK-mediated inhibition in either neuron subtype (Figure 5A,B). However, ACh release triggered increases in action potential frequency that were larger in CPn (n = 10) than in COM (n = 8) neurons (*p* = 0.006, one-tailed Student’s t-test), with mean increases in spike frequencies being 59 ± 13% in CPn neurons (*p* = 0.002, paired Student’s t-test) and 18 ± 4% (*p* = 0.001) in COM neurons (Figure 5C). Cholinergic excitation also persisted longer in CPn neurons (39 ± 10 s vs 14 ± 4 s in CPn and COM neurons, respectively; *p* = 0.028; one-tailed Student’s t-test; Figure 5C). Finally, when challenged with atropine (1 µM), flash-evoked excitation of CPn neurons was eliminated (from 67 ± 15% to 0 ± 2% above baseline firing rates; n = 8; p = 0.004, paired Student’s t-test; Figure 5D), confirming that mAChRs mediate responses to endogenous ACh in prefrontal projection neurons.

**Figure 5.**
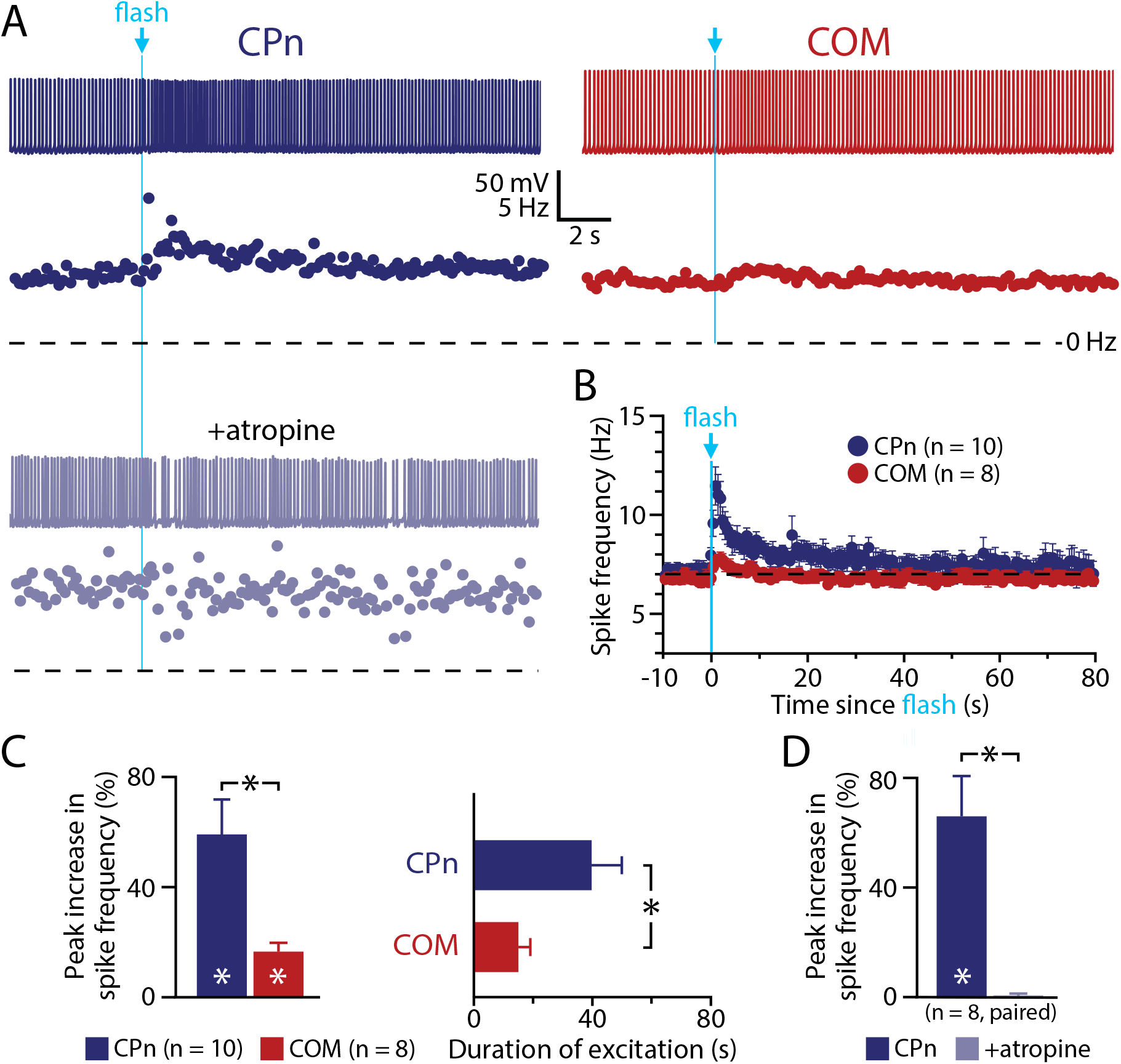
Preferential cholinergic excitation of CPn neurons in an alternative optogenetic model of endogenous ACh release. **A)** Voltage responses (top) and corresponding ISF plots (below) to single-flash-evoked release of endogenous ACh in CPn (blue trace, left) and COM (red trace, right) neurons in the mPFC of ChAT-Cre/Ai32 mice. In the CPn neuron, the flash-induced increase in firing frequency was eliminated following addition of atropine (1 µM; lower light blue trace and ISF plot). **B)** Plots of mean ISFs in during light-evoked release of endogenous ACh for populations of CPn (blue; n = 10) and COM (red; n = 8) neurons in the mPFC of ChAT-Cre / Ai32 mice. **C)** Comparisons of the magnitudes (left) and durations (right) of peak cholinergic excitation in CPn (n = 10) and COM (n = 8) neurons. **D)** Comparison of ACh response amplitudes in a subset of CPn neurons (n = 8) in baseline conditions and after application of atropine. Asterisks indicate significant differences (*p* < 0.05); white asterisks indicate significant changes from baseline firing frequencies, while black asterisks indicate significant differences between conditions.

### Mechanisms of persistent cholinergic excitation in CPn neurons

How does transient mAChR activation lead to persistent enhancement of CPn neuron excitability? To address this question, we first tested whether intrinsic properties of CPn neurons, or their connectivity within cortical networks, contribute to persistent excitation following a transient cholinergic stimulus. To rule out a role for intrinsic properties, we delivered a single 1 s current step to mimic transient enhanced excitatory drive in the middle of a longer suprathreshold current injection (Figure 6A). Across multiple current intensities, action potential frequencies increased during the period of added excitatory stimulus, but immediately (within 1.1 ± 0.1 ms; n = 3) returned to baseline or lower levels (Figure 6B), indicating that transient increases in firing frequency alone are not sufficient to drive long-lasting excitation of CPn neurons.

**Figure 6.**
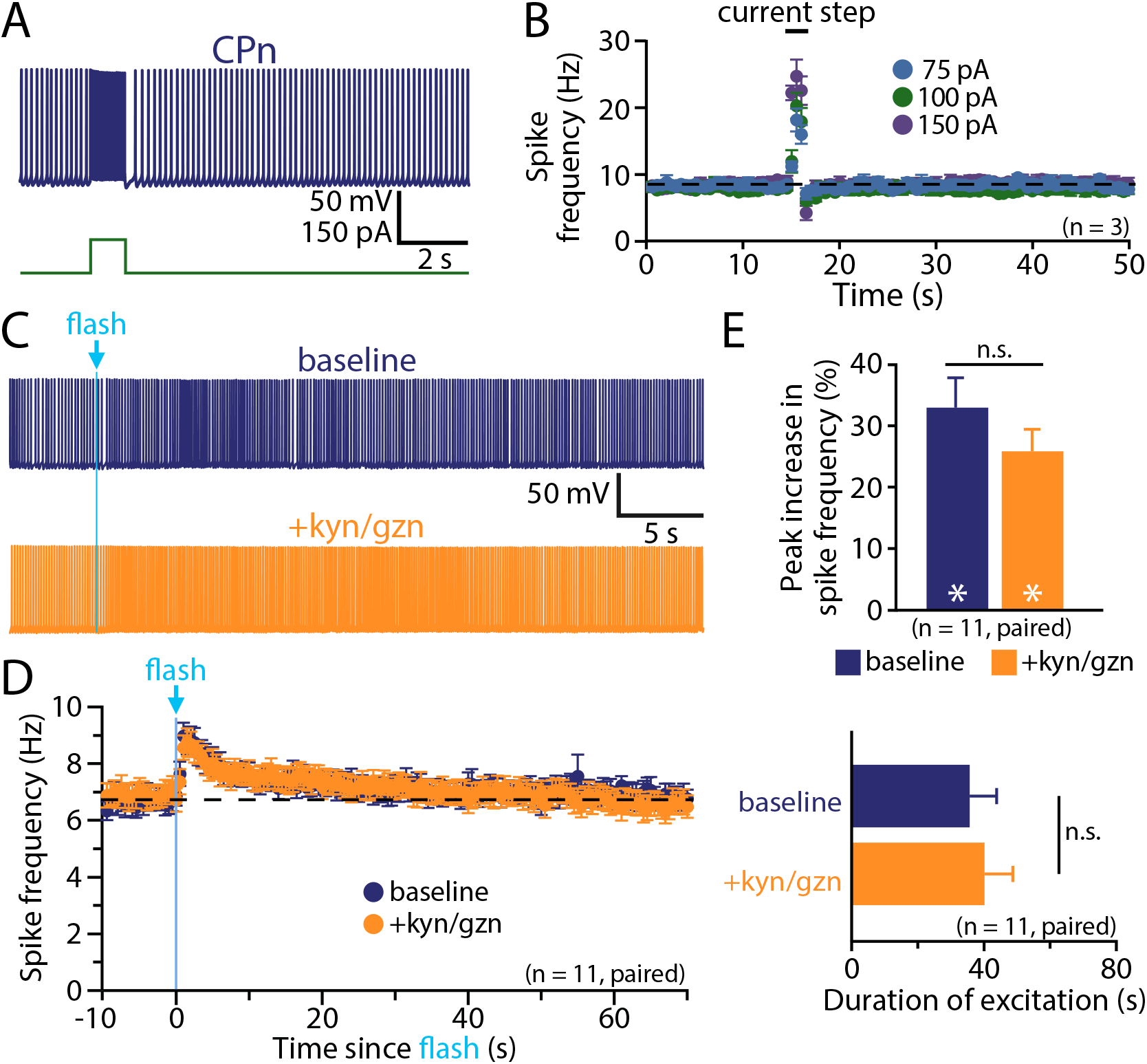
Persistence of cholinergic excitation of CPn neurons does not result from intrinsic membrane properties or network activity. **A)** Voltage response of a CPn neuron to a brief current step (100 pA, 1s) applied during spontaneous action potential generation driven by suprathreshold DC current. **B)** Plots of mean ISFs for CPn neurons (n = 3) experiencing transient increases in excitatory drive at the indicated intensities. Note that current-induced increases in spike frequency return to baseline levels immediately after cessation of the depolarizing step. **C)** Responses to flash-evoked release of endogenous ACh (5 ms) in a CPn neuron in baseline conditions (blue trace, top) and after block of fast synaptic transmission with kynurenate (kyn; 3 mM) and gabazine (gzn; 10 µM; orange trace, bottom). **D)** Plots of ISF for a population of CPn neurons (n = 11) in baseline conditions (blue) and in the presence of kyn and gzn (orange). **E)** Comparisons of the magnitude (top) or duration (bottom) of peak excitation for 11 CPn neurons in baseline conditions and after addition of kyn and gzn. White asterisks indicate significant differences (*p* < 0.05) from pre-flash firing frequencies.

To test the contribution of network activity in driving persistent cholinergic excitation, we measured responses of CPn neurons from ChAT-ChR2-YFP mice to single-flash-evoked endogenous ACh release before and after bath-application of kynurenate (3 mM), a nonspecific ionotropic glutamate receptor blocker, and gabazine (10 µM), a selective GABAa receptor blocker (Figure 6C,D). In baseline conditions, single-flash-induced increases in action potential output peaked at 33 ± 5% above baseline firing rates (*p* < 0.001, paired Student’s t-test; Figure 6E) and persisted for 36 ± 8 s (Figure 6F). After application of kynurenate and gabazine, flashes of light remained potent, enhancing action potent output by 26 ± 4% (*p* < 0.001; Figure 6E) and persisting for 40 ± 9 s (n = 11; Figure 6F). Thus, the magnitudes (*p* = 0.08; paired Student’s t-test) and durations (*p* = 0.63) of excitatory responses were similar in baseline and antagonist conditions, suggesting that changes in local network activity cannot account for persistent cholinergic excitation of CPn neurons.

We next tested ionic mechanisms that might contribute to cholinergic excitation of CPn neurons. Several forms of persistent cholinergic excitation require intracellular calcium signaling, including persistent firing triggered by nicotinic (Hedrick and Waters, 2015) and muscarinic (Egorov et al., 2002; Rahman and Berger, 2011) receptors in pyramidal neurons in primary sensorimotor cortices, as well as mAChR-dependent afterdepolarizations (ADPs) that occur following trains of action potentials in prefrontal pyramidal neurons (Dasari et al., 2013; Haj-Dahmane and Andrade, 1998; Yan et al., 2009). To determine whether persistent cholinergic excitation of CPn neurons requires intracellular calcium signaling, we chelated intracellular calcium by including BAPTA (10 or 30 mM) in patch pipettes. The efficacy of calcium chelation was confirmed by pairing exogenous ACh with brief spike trains to evoke calcium-dependent cholinergic ADPs. In control CPn neurons patched with normal pipette solution, transient applications of ACh paired with ten current-induced action potentials generated ADPs of 4.3 ± 0.7 mV (n = 5). These cholinergic ADPs were absent in neurons recorded with 10 mM (-0.4 ± 0.2 mV; n = 5) or 30 mM (0.0 ± 0.5 mV; n = 5) BAPTA in the pipette solution (Figure 7A), confirming successful chelation of internal calcium. Additionally, inclusion of BAPTA in the patch pipette eliminated SK-channel-dependent inhibitory responses to ACh (Figure 7B). However, while effective in blocking cholinergic ADPs and SK inhibition, inclusion of BAPTA in patch-pipettes failed to block persistent excitation by exogenous or endogenous ACh (Figure 7B,C). Instead, compared to cholinergic excitatory responses in control neurons (peak increases in ISF of 170 ± 8%, n = 57), focally applied ACh generated slightly larger excitatory responses (214 ± 15% above baseline spike frequencies; n = 47) in neurons filled with 10 mM BAPTA (p = 0.008; Student’s t-test), and equivalent excitation in neurons filled with 30 mM BAPTA (162 ± 28% above baseline frequencies, n = 15; p = 0.71; Figure 7D). Similarly, with intracellular calcium chelated with 10 mM BAPTA, excitatory responses to flash-evoked release of endogenous ACh were ~75% larger (mean increase in ISF = 47 ± 5%; n = 27) than responses in control neurons (mean increase = 27 ± 3%; n = 27, p = 0.002 vs control neurons, Student’s t-test; Figure 7D). Chelation of internal calcium also failed to reduce the persistence of cholinergic excitation, with durations of responses to exogenous ACh (59 ± 4 s for 10 mM BAPTA; p = 0.20 vs control; 43 ± 8 s for 30 mM BAPTA; p = 0.24 vs control) or endogenous ACh (31 ± 6 s for 10 mM BAPTA; p = 0.37 vs control) being similar to the durations of responses in control neurons lacking internal BAPTA (exogenous, 52 ± 3 s, n = 57; endogenous, 23 ± 5 s, n = 27; Figure 7D). Thus, chelation of intracellular calcium failed to reduce the magnitudes or durations of cholinergic excitatory responses in CPn neurons, even as it blocked fully any contribution of the ADP conductance. Instead, 10 mM BAPTA enhanced the magnitude of excitation in response to both exogenous and endogenous ACh. These results demonstrate that intracellular calcium signaling is not necessary for cholinergic enhancement of action potential output in CPn neurons, and suggest instead that the net effect of normal intracellular calcium levels may be to suppress cholinergic responses.

**Figure 7.**
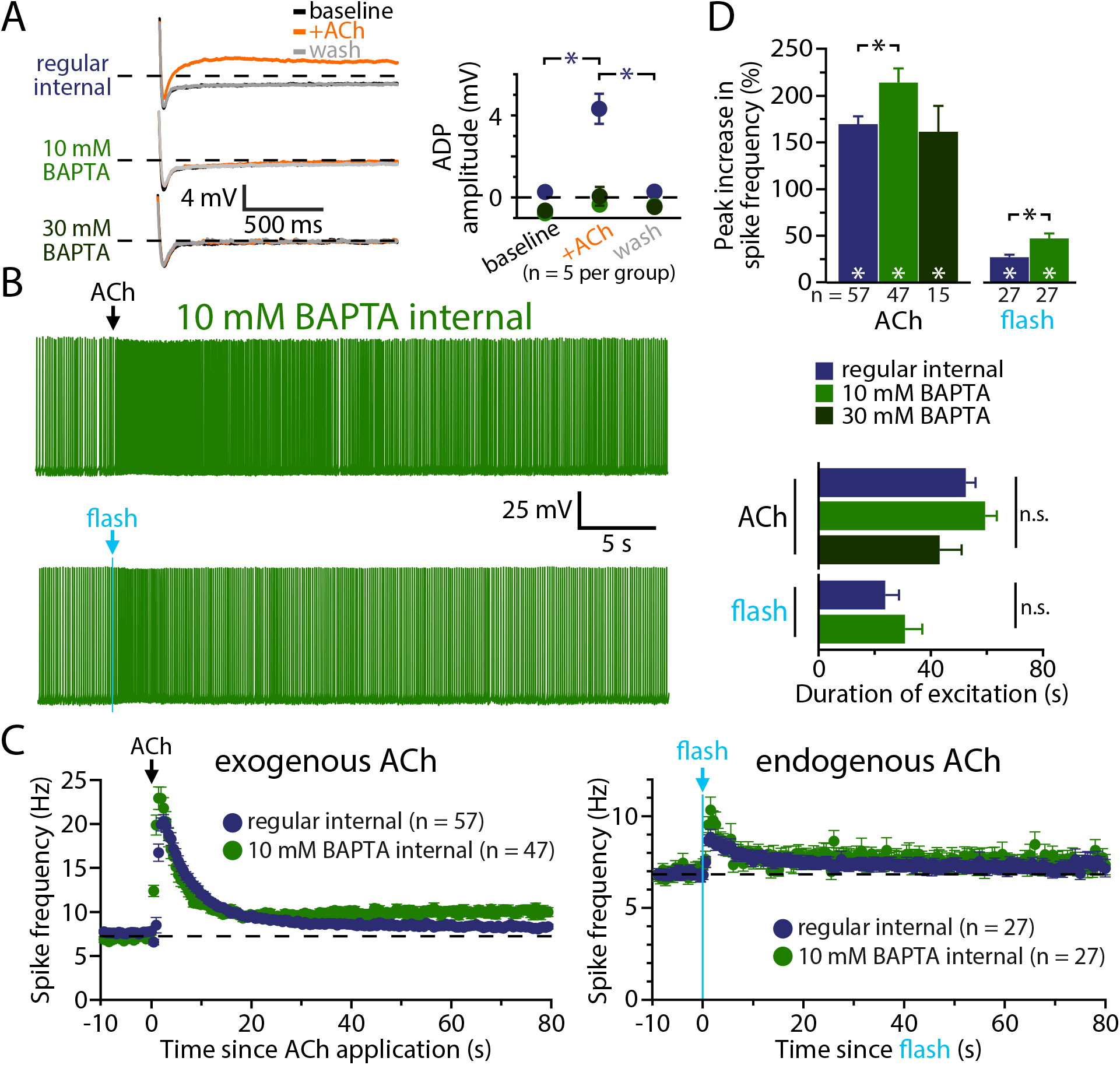
Chelating internal calcium does not reduce cholinergic excitatory responses in CPn neurons. **A)** Pairing three applications of exogenous ACh with ten current-driven action potentials led to the generation of afterdepolarizing potentials (ADPs) in a CPn neuron recorded with control internal (top traces), but not in neurons filled with 10 mM (middle traces) or 30 mM (bottom traces) BAPTA (left). Quantification of ADP amplitudes before, during, and after exposure to ACh in control neurons (blue; n = 5) and neurons filled with 10 mM (light green; n = 5) or 30 mM (dark green; n = 5) BAPTA (right). **B)** Voltage responses to exogenous (top) and endogenous (bottom) ACh in neurons filled with 10 mM BAPTA. **C)** Plots of mean ISFs for populations of CPn neurons in response to exogenous (left) or endogenous (right) ACh recorded with regular internal (blue), 10 mM BAPTA internal (light green), or 30 mM BAPTA internal (dark green). **D)** Comparisons of the peak increase in firing frequency (top) or duration of excitation (bottom) in response to exogenous or endogenous ACh in control neurons and neurons filled with 10 or 30 mM BAPTA. Asterisks indicate significant differences (*p* < 0.05); white asterisks indicate significant changes from baseline firing frequencies, while black asterisks indicate significant differences between conditions.

One calcium-sensitive ionic mechanism classically associated with Gq-coupled receptors is the M-current (Delmas et al., 2004). To test the role of M-current in cholinergic excitation, we measured responses to focal application of exogenous ACh in baseline conditions (control internal, left, or BAPTA internal, right) and after bath-application of XE991 (10 µM), a selective blocker of the Kv7 potassium channels that underly the M-current (Figure 8A,B). With internal calcium intact, addition of XE991 reduced peak excitatory responses by 23 ± 9% (n = 11; *p* = 0.018, repeated measures ANOVA), from 197 ± 18% increases in ISF in baseline conditions to 142 ± 13% increases in XE991 (Figure 8C). Similarly, in a second set of neurons filled with 10 mM BAPTA, addition of XE991 reduced peak excitatory responses by 19 ± 5% (n = 16; *p* < 0.001), from 219 ± 30% increases in spike frequencies in baseline conditions to 174 ± 26% increases in XE991 (Figure 8C). XE991 also reduced the persistence of cholinergic excitatory responses by 29 ± 24% (from 53 ± 8 s to 28 ± 7 s) with internal calcium intact (p = 0.03), and by 24 ± 10% (from 56 ± 7 s to 42 ± 9 s) with internal calcium chelated with BAPTA (p = 0.06; Figure 8C). The effects of XE991 on cholinergic excitatory responses did not wash out within 15 minutes. These findings suggest that suppression of the M-current contributes to persistent cholinergic excitation of CPn neurons. However, because the efficacy of XE991 was not enhanced in BAPTA-filled neurons, it is unlikely that calcium-sensitivity of M-current (Selyanko and Brown, 1996) accounts for the larger response amplitudes observed in the presence of BAPTA.

**Figure 8.**
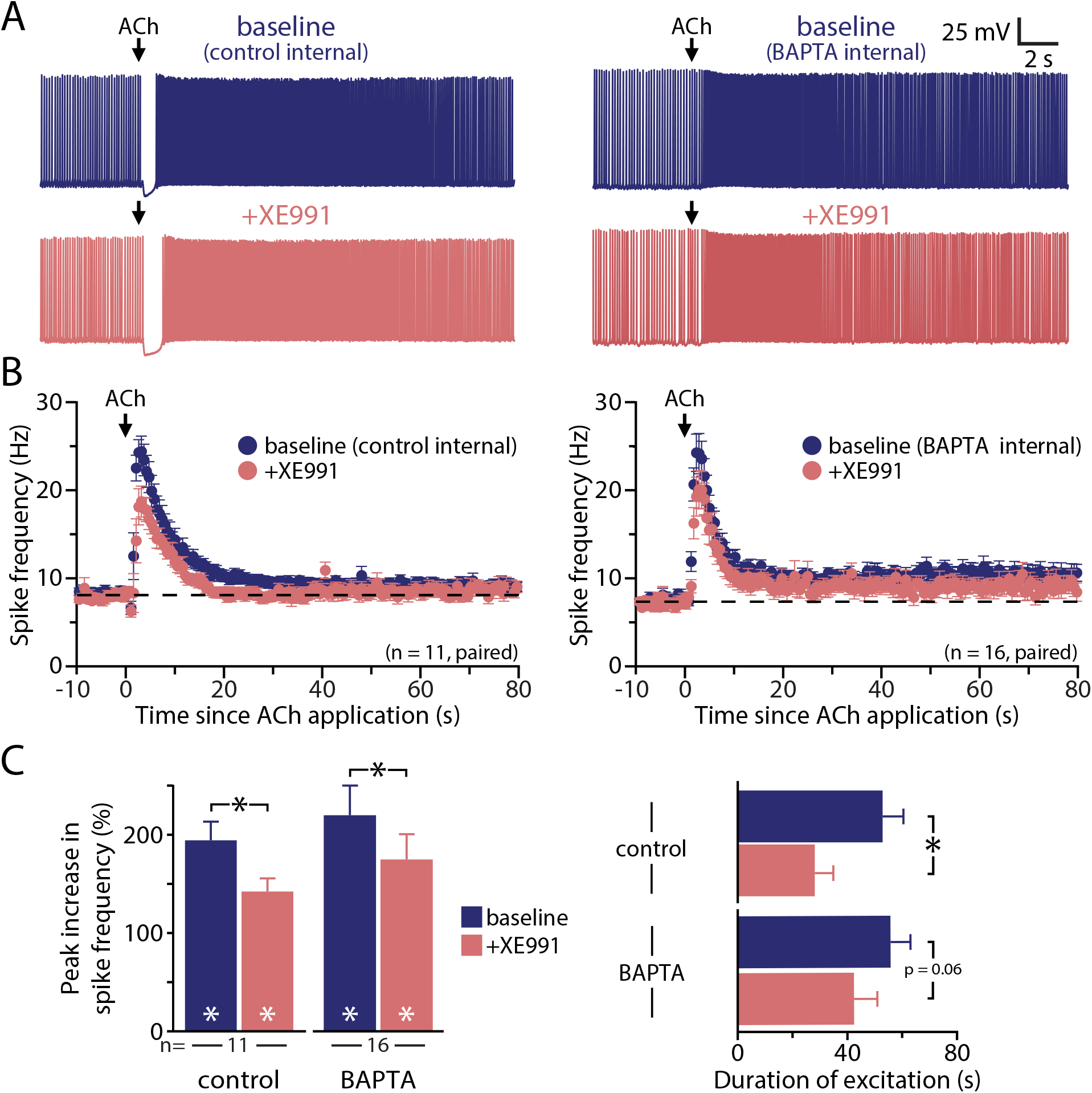
M-current contributes to persistent cholinergic excitation of CPn neurons. **A)** Voltage responses to focal application of exogenous ACh (100 ms) in a CPn neuron recorded with control internal (left) or 10 mM BAPTA internal (right) in baseline conditions (top, blue) and after addition of the Kv7 blocker XE911 (10 µM; bottom, pink). **B)** Plot of mean ISFs for a population of CPn neurons (control internal, n = 11, left; 10 mM BAPTA internal, n = 16, right) in baseline conditions (blue) and after blockade of Kv7 channels with XE991 (pink). **C)** Comparisons of the magnitude (left) and durations (right) of cholinergic responses in baseline conditions and after addition of XE991 in control and BAPTA-filled CPn neurons. Asterisks indicate significant differences (*p* < 0.05); white asterisks indicate significant changes from pre-ACh firing frequencies, while black asterisks indicate significant differences between conditions.

While intracellular calcium signaling may not be required for cholinergic excitation of CPn neurons, mAChR activation promotes calcium influx (Dasari et al., 2017; Power and Sah, 2005). To test whether calcium conductances contribute to cholinergic excitation, we used two approaches: blockade of calcium conductances with bath applied cadmium (200 µM), and removal of extracellular calcium. Because these manipulations block synaptic transmission, we used focal applications of exogenous ACh in these experiments. We first measured ACh responses in CPn neurons filled with either control intracellular solution, or a solution containing 10 mM BAPTA, before and after addition of cadmium to the aCSF (Figure 9A). In control conditions (i.e., in the absence of intracellular BAPTA), application of cadmium did not change mean response amplitudes (a 140 ± 18% increase in ISF in baseline conditions vs a 152 ± 26% in the presence of cadmium; n = 14, *p* = 0.610, paired Student’s t-test). However, cadmium application markedly reduced the duration of cholinergic excitation by 67 ± 7%, from 56 ± 7 s to 14 ± 1 s (n = 14; p < 0.001, Student’s paired t-test). With intracellular calcium chelated with BAPTA (n = 10), baseline ACh-induced increases in ISF (240 ± 28%) were larger than in control neurons (P = 0.005 vs control neurons lacking BAPTA). Application of cadmium to BAPTA-filled CPn neurons led to a 47 ± 8% reduction in response amplitudes (mean increase of ISF in cadmium was 135 ± 26%; p < 0.001; Figure 9C). Cadmium also reduced the duration of cholinergic responses in BAPTA-filled neurons by 59 ± 10%, from 54 ± 11 s to 20 ± 6 s (p = 0.003). These results suggest ACh activates a calcium-sensitive calcium conductance that contributes to the amplitude and persistence of cholinergic excitatory responses in CPn neurons, and confirm that intracellular calcium signaling has a net effect of suppressing cholinergic excitation.

**Figure 9.**
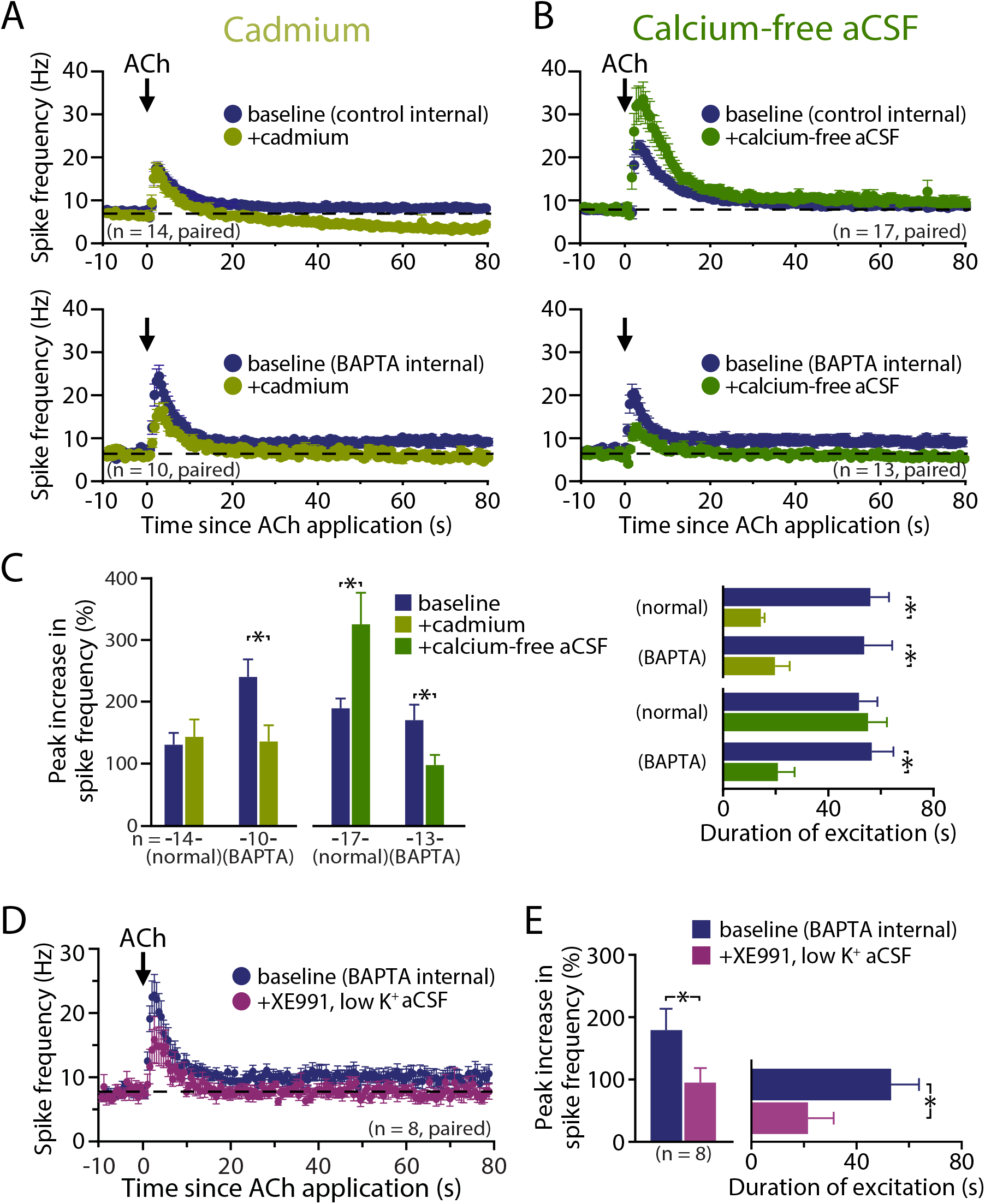
Blockade of inward calcium conductances reduces persistence of cholinergic excitation. **A, B)** Plots of mean ISFs over time in populations of neurons exposed to exogenous ACh in baseline (blue) or after manipulation of calcium conductances (dark green, addition of 200 µM cadmium, **A**; light green, transition to calcium-free aSCF, **B)**. **C)** Comparisons of the peak increases in firing frequency (left) and response durations (right) in CPn neurons recorded with normal (i.e., non-BAPTA) intracellular solution, or with 10 mM intracellular BAPTA, before and after addition of cadmium (light green) or removal of external calcium (dark green). **D)** Plots of mean ISFs for a population of CPn neurons in baseline conditions (blue) and after adding XE991 (10 µM) in low potassium (0.5 mM) aCSF (purple; n = 8). **E)** Comparisons of the magnitude (left) and duration (right) of excitatory responses in baseline (BAPTA) conditions and in the presence of XE991 and 0.5 mM external potassium. Asterisks indicate significant differences (*p* < 0.05) between conditions.

To further test the role of calcium conductances in mediating cholinergic excitation, we exposed control and BAPTA-filled CPn neurons to ACh before and after exchanging our normal aCSF with nominally calcium-free aCSF containing 3 mM Mg^2+^ (Figure 9B). With internal calcium intact, transition to calcium-free aCSF enhanced the magnitude of cholinergic responses by 66 ± 22%, with peak increases in ISF moving from 190 ± 16% in baseline conditions to 325 ± 51% above initial firing rates after removal of extracellular calcium (n = 17, *p* = 0.006, repeated measures ANOVA; Figure 9C), an effect that was reversible (to 141 ± 21%, p < 0.001) upon reintroduction of external calcium. Removal of external calcium did not affect the persistence of cholinergic excitation, with durations of enhanced action potential output being similar (52 ± 7 s in baseline conditions vs 55 ± 7 s in the absence of external calcium; p = 0.66; Figure 9C). On the other hand, when intracellular calcium was chelated with BAPTA (10 mM), calcium-free aCSF reduced peak increases in ISF by 42 ± 8%, from 169 ± 25% to 97 ± 16% (n = 13, *p* = 0.002, repeated measures ANOVA; Figure 9C), an effect that was partially reversible (to 136 ± 18%; p = 0.032) upon reintroduction of external calcium. Removal of external calcium also reversibly reduced the persistence of cholinergic excitation in BAPTA-filled neurons by 49 ± 20%, with durations of enhanced action potential output dropping from 56 ± 8 s in baseline conditions to 21 ± 6 s in the absence of external calcium (*p* = 0.005), but returning to 46 ± 9 s after reintroduction of calcium (*p* = 0.049 vs calcium-free condition; Figure 9C). These results further suggest that cholinergic excitation of CPn neurons involves activation of a calcium-permeable nonspecific cation conductance that under normal conditions is negatively regulated by intracellular calcium.

Finally, to test whether this remaining calcium conductance represents a nonspecific cation conductance, we measured responses to exogenous ACh in BAPTA-filled CPn neurons before and after application of XE991 (10 µM) in “low-K^+^” aCSF containing 0.5 mM potassium. This treatment, which enhances the driving force for potassium while also blocking the M-current, reduced peak excitation by 46 ± 10% (from 179 ± 34% to 95 ± 23%; n = 8, *p* = 0.007; Figure 9D), and response persistence by 44 ± 26% (from 53 ± 11 s to +21 ± 9 s; p = 0.06; Figure 9E). The suppressive effect of XE991 and low extracellular potassium on response amplitude in BAPTA-filled CPn neurons was significantly greater than that of XE991 alone (n = 24; p = 0.011, Student’s t-test), suggesting that the remaining calcium-permeable conductance (i.e., after blockade of K_v_7 channels and ADP channels with XE991 and BAPTA, respectively) is a nonspecific cation channel with potassium permeability. However, the two treatments (XE991 alone and XE991 with reduced extracellular potassium) had similar effects on the duration of cholinergic responses (p = 0.408), indicating that this conductance may be primarily involved in the early part of the response, or that cholinergic suppression of other, non-K_v_7 potassium conductances contribute selectively to response persistence.

Based on our findings above that cholinergic excitation involves both suppression of K_v_7 conductances and activation of nonspecific cation conductances, we next tested for interaction of these mechanisms by challenging cholinergic responses to combined elimination of both components. We found that co-application of XE991 and cadmium triggered spontaneous up-down states in neurons (Figure 10A), making it impossible to establish consistent baseline firing frequencies from which to measure cholinergic responses. However, since blockade of calcium conductances and removal of extracellular calcium resulted in qualitatively similar reductions in cholinergic responses in BAPTA-filled neurons, we measured responses to exogenous ACh in CPn neurons before and after co-applying XE991 (10 µM) and calcium-free aCSF in BAPTA-filled CPn neurons (Figure 10B,C). Addition of XE991 in calcium-free aCSF reduced the magnitude of cholinergic excitatory responses by 71 ± 6% (from 258 ± 38% to 74 ± 19% increases in ISF; n = 11, *p* < 0.001, repeated measures ANOVA), an effect that was partially reversible (to 123 ± 14%; p = 0.003) after adding back external calcium and removing XE991. Application of calcium-free aCSF and XE991 also reduced the duration of excitation by 78 ± 9% (from 75 ± 5 s to 15 ± 7 s; n = 11, *p* < 0.001), and this reduction was partially rescued by reintroduction of extracellular calcium (to 53 ± 8 s, p = 0.007; Figure 10B,C), consistent with our previous findings that the impact of calcium removal, but not XE991, is reversible.

**Figure 10.**
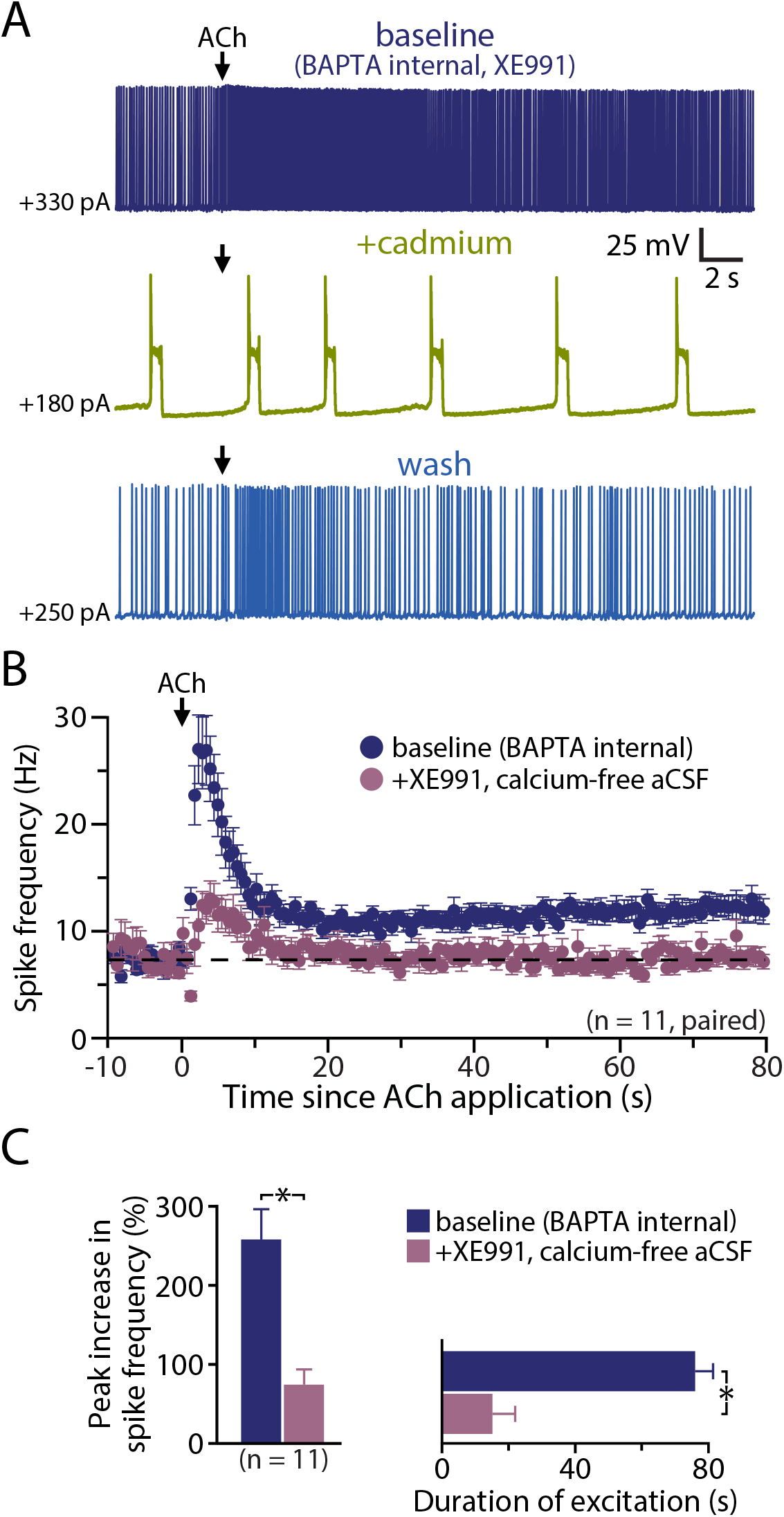
Cholinergic excitation involves M-current and a calcium-conductance. **A)** Voltage response of a CPn neuron to ACh (100 ms) in baseline conditions (10 mM BAPTA internal and XE991, dark blue trace), after addition of cadmium (200 µM, green trace), and during wash (light blue trace). Note presence of up-down states upon combination of BAPTA, XE991, and cadmium. **B)** Plots of mean ISFs for a population of CPn neurons in baseline conditions and after transition to XE991 (10 µM) in calcium-free aCSF (n = 11). **C)** Comparisons of the magnitude (left) and duration (right) of excitatory responses in baseline (BAPTA) conditions (blue) and in the presence of XE991 and calcium-free aCSF (n = 11). Asterisks indicate significant differences (*p* < 0.05) between conditions.

Together, these findings suggest that the M-current and a calcium-permeable nonspecific conductance independently and additively contribute to cholinergic excitation of CPn neurons. An additional ionic contributor under normal conditions likely includes the calcium-gated nonspecific cation conductance underlying the ADP, which is activated when M1 receptor activation is paired with action potential generation (Haj-Dahmane and Andrade, 1998). However, our finding that chelation of intracellular calcium did not reduce the amplitude of cholinergic excitation suggests that the contribution of the ADP current to excitatory responses must be balanced by calcium-dependent suppression of other conductances (see Discussion, below).

## Discussion

A growing appreciation of the diversity of cortical neurons, their selective connectivity, and their differential responsivity to neuromodulatory transmitters is expanding our understanding of cortical circuit function (Dembrow and Johnston, 2014). While pyramidal neurons broadly express postsynaptic M1-subtype mAChRs and are excited by muscarinic agonists (Dasari and Gulledge, 2011; Gulledge et al., 2009), recent studies have revealed projection-specific cholinergic signaling in a variety of cortical areas, including prefrontal (Dembrow et al., 2010), motor (Hedrick and Waters, 2015), and auditory (Joshi et al., 2016) cortices. Our main finding is that transient release of endogenous ACh, as occurs in the mPFC during cue-detection tasks (Parikh et al., 2007), selectively and persistently enhances action potential generation in prefrontal CPn neurons. Cholinergic excitation was robust, occurring after even a single ACh-release event, lasted for many tens of seconds, and was mediated by a combination of ionic effectors, including K_v_7 and at least two nonspecific cation conductances. As discussed below, preferential cholinergic excitation of CPn neurons may contribute to circuit-based mechanisms in the mPFC that facilitate attentional processes.

### Cholinergic responses in cortical projection neurons

Consistent with broad expression of M1 receptors in pyramidal neurons (Dasari and Gulledge, 2011; Gulledge et al., 2009), exogenous ACh generated qualitatively similar responses, including transient inhibition followed by longer-lasting excitation, in CPn and COM neurons in the mPFC. Yet, the overall impact of exogenous ACh on excitability was projection-specific, as COM neurons had longer lasting apamin-sensitive inhibitory responses, while CPn neurons exhibited larger, and longer lasting, excitation. Preferential excitation of CPn neurons was not due to interaction of SK-mediated inhibition, as excitatory responses to exogenous ACh remained larger in CPn neurons when SK channels were blocked, and endogenous ACh release in two different optogenetic mouse models preferentially excited CPn neurons in the absence of cholinergic inhibition. These results are consistent with those of Dembrow et al. (2010), who found that tonic mAChR activation with bath-applied agonists preferentially enhanced the excitability of CPn neurons, and suggest that transient release of ACh in the mPFC, as occurs during cue-detection tasks (e.g., Parikh et al., 2007), will also preferentially facilitate corticofugal output from the PFC (Figure 11A).

**Figure 11.**
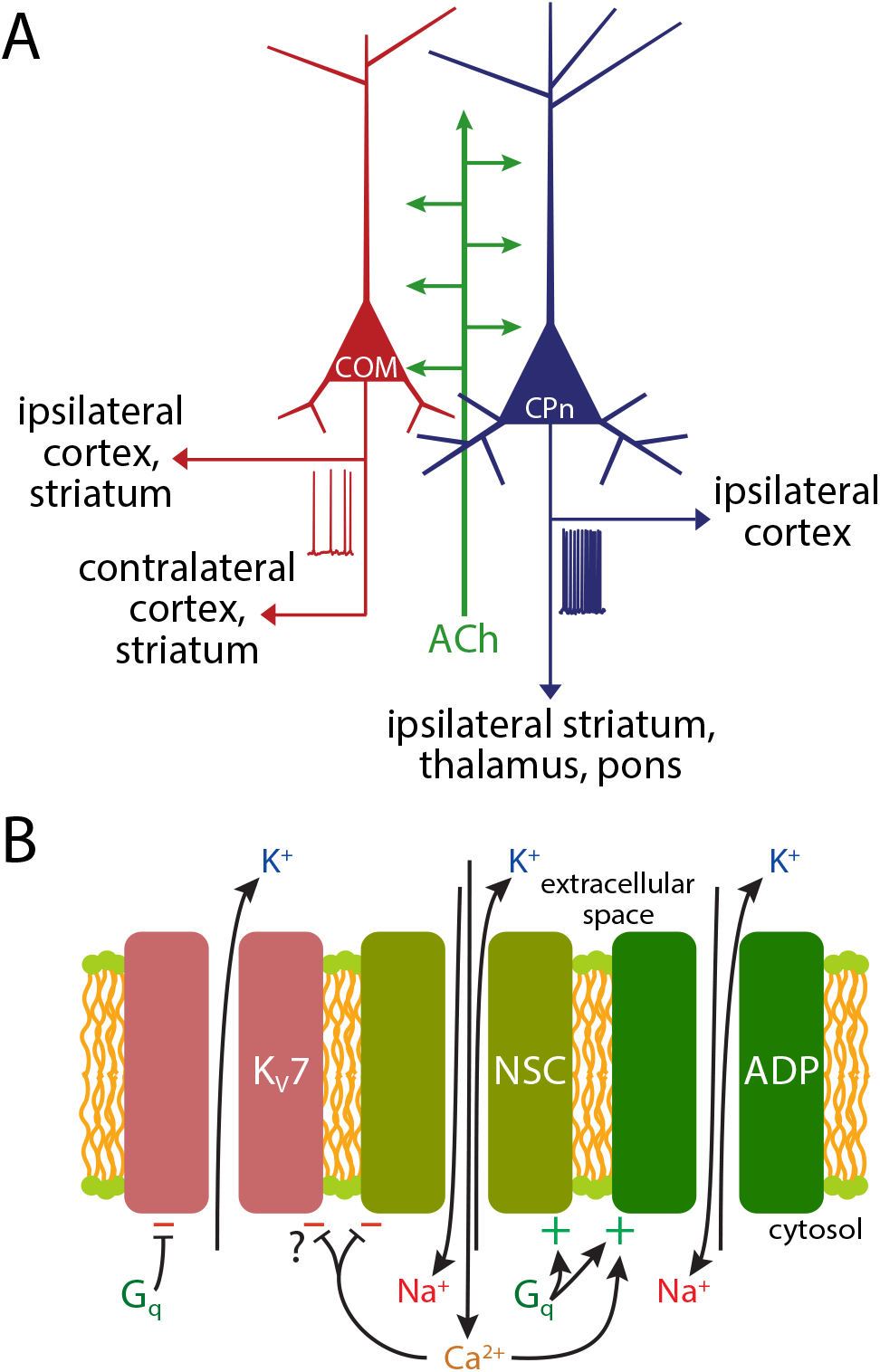
Model of cholinergic regulation of cortical projection neurons. **A)** Diagram of cholinergic regulation of cortical projection neurons neurons. ACh promotes cortical output to the brainstem by preferentially exciting corticopontine (CPn) neurons relative to commissural/callosal (COM) neurons. **B)** Diagram of three ionic mechanisms contributing to Gq-triggered excitation in CPn neurons: suppression of Kv7 channels (M-current), activation of the nonspecific cation conductance underlying the calcium-dependent afterdepolarization (ADP), and activation of a calcium-sensitive but calcium-permeable nonspecific cation (NSC) conductance.

In layer 5 neurons of the motor (Hedrick and Waters, 2015) and auditory (Joshi et al., 2016) cortices, endogenous ACh release activates both mAChRs and nicotinic acetylcholine receptors (nAChRs) to generate a sundry of postsynaptic responses in pyramidal neurons, ranging from mAChR-mediated inhibition to fast or slow depolarizations mediated by nAChRs and/or mAChRs. We did not observe nAChR-mediated responses in layer 5 pyramidal neurons in the mPFC (see also Porter et al., 1999), and responses to both exogenous and endogenous ACh were eliminated in the presence of the muscarinic antagonist atropine. This likely reflects intrinsic differences in nAChR expression in layer 5 neurons across cortical areas (Hedrick and Waters, 2015). Similarly, we did not observe significant SK-mediated inhibition in either neuron subtype following endogenous release of ACh. However, SK responses to exogenous ACh were larger in COM neurons, a result consistent with the preferential cholinergic inhibition of COM neurons observed in the auditory cortex (Joshi et al., 2016). Because inhibitory responses are robust in COM neurons across cortical areas, and both inhibitory and excitatory responses rely on M1 receptors (Gulledge et al., 2009), it is likely that cell-type specificity in cholinergic actions reflect differences in intracellular signaling cascades and/or subcellular localization of M1 receptors, rather than quantitative differences in receptor expression or cholinergic innervation patterns.

Our first optogenetic animal model, the ChAT-ChR2-YFP mouse, has the disadvantage of overexpressing the vesicular ACh transporter (VAChT), potentially leading to larger-than-normal quantal content in cholinergic vesicles and enhanced “cholinergic tone” (Kolisnyk et al., 2013b). Our second model, a cross between ChAT-IRES-Cre mice expressing Cre in cholinergic neurons, and Ai32 mice driving Cre-dependent expression of channelrhodopsin-2, has normal VAChT expression, but may drive ChR2 expression in non-cholinergic neurons in a minority of mice (Hedrick et al., 2016). Our results were consistent across both animal models in revealing preferential excitation of prefrontal CPn neurons by endogenous ACh. Since responses were eliminated by atropine, but not by blockers of glutamate and GABA receptors, they cannot be attributed to co-transmission of glutamate (Allen et al., 2006) or optical activation of non-cholinergic neurons (Hedrick et al., 2016), but instead appear to result from activation of postsynaptic mAChRs. This conclusion is consistent with results from both Dembrow et al. (2010) and Joshi et al. (2016), and suggests a generalized role for muscarinic receptor signaling in amplifying corticofugal output throughout the neocortex.

### Persistent cholinergic excitation of CPn neurons

Tonic activation of mAChRs promotes persistent spontaneous action potential generation in pyramidal neurons (Andrade, 1991; Dembrow et al., 2010; Egorov et al., 2002; Gulledge et al., 2009; Rahman and Berger, 2011). Similarly, transient release of endogenous ACh in motor (Hedrick and Waters, 2015) and sensory (Joshi et al., 2016) cortices promotes persistent action potential generation when neurons are depolarized close to, or beyond, action potential threshold. Our results from experiments using two different optogenetic models for endogenous ACh release are consistent with those of Joshi et al. (2016) in that persistent cholinergic excitation depended purely on mAChR activation, lasted for many tens of seconds, and occurred preferentially in corticofugal projection neurons.

At least three distinct mechanisms likely contribute to persistent muscarinic excitation of CPn neurons in the mPFC: inhibition of Kv7 channels mediating the M-current and activation of two nonspecific cation conductances, including the calcium-gated conductance underlying the ADP and another calcium-permeable nonspecific cation conductance (Figure 11B). Cholinergic ADPs following bursts of action potentials are gated by calcium influx during spike trains (Andrade, 1991; Dasari et al., 2013; Haj-Dahmane and Andrade, 1998; Yan et al., 2009), and therefore likely contribute to sustaining cholinergic excitation during ongoing action potential genesis. Yet, we found muscarinic excitation of layer 5 neurons to be unaffected, or enhanced, after chelation of intracellular calcium (see Figure 7). Such calcium-independent muscarinic excitation has been observed in pyramidal neurons in several cortical areas (Egorov et al., 2003; Guerineau et al., 1995; Haj-Dahmane and Andrade, 1996; Shalinsky et al., 2002) and likely reflects a combination of cholinergic inhibition of Kv7 channels and activation of an additional calcium-permeable nonspecific cation conductance that may itself be inhibited by intracellular calcium (Magistretti et al., 2004). Unfortunately, we were not able to determine the relative contributions of the ADP and calcium-permeable conductances in control conditions, as manipulations to one conductance also affected the other. For instance, inclusion of intracellular BAPTA to block the ADP also enhanced the calcium-permeable conductance. Similarly, addition of cadmium or removal of extracellular calcium to manipulate the calcium conductance also eliminated the calcium-dependent ADP. One intriguing idea is that these two conductances are balanced such that cholinergic excitation remains robust regardless of the state of intracellular calcium. This would explain the modest effect of intracellular BAPTA on response amplitudes. Further, the intrinsic negative feedback of the calcium-sensitive calcium conductance provides a mechanism to stabilize intracellular calcium levels, which may contribute to calcium store refilling after Gq-triggered calcium release events (Dasari et al., 2017).

With intracellular calcium signaling blocked with BAPTA (i.e., in the absence of the calcium-gated ADP conductance and SK-channel-mediated inhibition), we found cholinergic excitation of CPn neurons to be independently sensitive to blockade of K_v_7 channels (~20 to 25% reductions in amplitudes and durations), and blockade of calcium conductances with cadmium (~50 to 60% reductions) or removal of external calcium (~40 to 50% reductions). These effects were additive, such that the combined removal of calcium and addition of XE991 generated a ~70% decrease in peak cholinergic excitation and ~80% reduction in response duration when BAPTA was included in patch pipettes. The residual excitation occurring in the absence of M-current and extracellular calcium may result from incomplete blockade of channels, or the presence of additional ionic effectors.

### Functional implications of selective cholinergic excitation of CPn neurons

Two forms of attention, stimulus-driven (“bottom-up”) and goal-oriented (“top-down”) attention, are differentiated based on whether attention is triggered by salient external stimuli (bottom-up) or by internal expectation of stimuli (top-down; Pinto et al., 2013). Although muscarinic signaling in the cortex is broadly associated with top-down attentional mechanisms (Herrero et al., 2008; McGaughy et al., 2002; Newman and McGaughy, 2008), transient rises of ACh in the mPFC gate bottom-up attention during cue detection tasks in which rodents orient toward salient stimuli predicting food reward (Gritton et al., 2016; Parikh et al., 2007). As CPn output drives cortical-cerebellar motor circuits, it is likely that cholinergic activation of CPn neurons during cue-detection tasks contributes to initiation of cue-evoked behavior. Consistent with this, the timing of cholinergic transients in the mPFC of rats is highly correlated with the initiation of cue-evoked behaviors (Parikh et al., 2007). More speculatively, it is possible that cholinergic excitation of prefrontal CPn neurons also acts to couple bottom-up and top-down attentional mechanisms in two ways. First, axon collaterals of layer 5 pyramidal neurons targeting the basal forebrain may promote ACh release in other cortical areas (Gielow and Zaborszky, 2017), as activation of the PFC, including via mAChR stimulation, is sufficient to induce ACh release in the parietal cortex (Nelson et al., 2005) and is necessary for sensory-evoked ACh release in primary sensory areas (Rasmusson et al., 2007). Second, intracortical CPn axon collaterals contribute to top-down feedback projections within cortical hierarchies (Ueta et al., 2013; Ueta et al., 2014) to provide prospective information to lower order cortical areas (Larkum, 2013). Thus, cue-driven muscarinic activation of prefrontal CPn neurons may promote top-down attentional processing by stimulating cortical release of ACh in relevant cortical target areas while also providing feedback corticocortical glutamatergic drive to guide attention toward relevant environmental stimuli. These hypotheses are testable by identifying the subtype(s) of cortical neurons that innervate cholinergic neurons in the basal forebrain, and by manipulating feedback corticocortical communication during attentional tasks *in vivo*.

## References

Allen, T.G., Abogadie, F.C., and Brown, D.A. (2006). Simultaneous release of glutamate and acetylcholine from single magnocellular “cholinergic” basal forebrain neurons. The Journal of neuroscience: the official journal of the Society for Neuroscience 26, 1588–1595.

Andrade R. (1991). Cell excitation enhances muscarinic cholinergic responses in rat association cortex. Brain Research 548, 81–93.

Bentley, P., Husain, M., and Dolan, R.J. (2004). Effects of cholinergic enhancement on visual stimulation, spatial attention, and spatial working memory. Neuron 41, 969–982.

Dalley, J.W., Theobald, D.E., Bouger, P., Chudasama, Y., Cardinal, R.N., and Robbins, T.W. (2004). Cortical cholinergic function and deficits in visual attentional performance in rats following 192 IgG-saporin-induced lesions of the medial prefrontal cortex. Cereb Cortex 14, 922–932.

Dasari, S., Abramowitz, J., Birnbaumer, L., and Gulledge, A.T. (2013). Do canonical transient receptor potential channels mediate cholinergic excitation of cortical pyramidal neurons? Neuroreport 24, 550–554.

Dasari, S., and Gulledge, A.T. (2011). M1 and M4 receptors modulate hippocampal pyramidal neurons. J Neurophysiol 105, 779–792.

Dasari, S., Hill, C., and Gulledge, A.T. (2017). A unifying hypothesis for M1 muscarinic receptor signalling in pyramidal neurons. The Journal of Physiology 595, 1711–1723.

Delmas, P., Padilla, F., Osorio, N., Coste, B., Raoux, M., and Crest, M. (2004). Polycystins, calcium signaling, and human diseases. Biochemical and biophysical research communications 322, 1374–1383.

Dembrow, N., and Johnston, D. (2014). Subcircuit-specific neuromodulation in the prefrontal cortex. Frontiers in neural circuits 8, 54.

Dembrow, N.C., Chitwood, R.A., and Johnston, D. (2010). Projection-specific neuromodulation of medial prefrontal cortex neurons. J Neurosci 30, 16922–16937.

Egorov, A.V, Angelova, P.R., Heinemann, U., and Müller, W. (2003). Ca2+-independent muscarinic excitation of rat medial entorhinal cortex layer V neurons. The European journal of neuroscience 18, 3343–3351.

Egorov, A.V, Hamam, B.N., Fransen, E., Hasselmo, M.E., and Alonso, A.A. (2002). Graded persistent activity in entorhinal cortex neurons. Nature 420, 173–178.

Gee, S., Ellwood, I., Patel, T., Luongo, F., Deisseroth, K., and Sohal, VS. (2012). Synaptic activity unmasks dopamine D2 receptor modulation of a specific class of layer V pyramidal neurons in prefrontal cortex. J Neurosci 32, 4959–4971.

Gielow, M.R., and Zaborszky, L. (2017). The Input-Output Relationship of the Cholinergic Basal Forebrain. Cell Rep 18, 1817–1830.

Gritton, H.J., Howe, W.M., Mallory, C.S., Hetrick, V.L., Berke, J.D., and Sarter, M. (2016). Cortical cholinergic signaling controls the detection of cues. Proc Natl Acad Sci U S A 113, E1089–1097.

Guerineau, N.C., Bossu, J.L., Gahwiler, B.H., and Gerber, U. (1995). Activation of a nonselective cationic conductance by metabotropic glutamatergic and muscarinic agonists in CA3 pyramidal neurons of the rat hippocampus. J Neurosci 15, 4395–4407.

Gulledge, A.T., Bucci, D.J., Zhang, S.S., Matsui, M., and Yeh, H.H. (2009). M1 receptors mediate cholinergic modulation of excitability in neocortical pyramidal neurons. J Neurosci 29, 9888–9902.

Gulledge, A.T., and Stuart, G.J. (2005). Cholinergic inhibition of neocortical pyramidal neurons. Journal of Neuroscience 25, 10308–10320.

Haj-Dahmane, S., and Andrade, R. (1996). Muscarinic activation of a voltage-dependent cation nonselective current in rat association cortex. J Neurosci 16, 3848–3861.

Haj-Dahmane, S., and Andrade, R. (1998). Ionic mechanism of the slow afterdepolarization induced by muscarinic receptor activation in rat prefrontal cortex. J Neurophysiol 80, 1197–1210.

Hedrick, T., Danskin, B., Larsen, R.S., Ollerenshaw, D., Groblewski, P., Valley, M., Olsen, S., and Waters, J. (2016). Characterization of Channelrhodopsin and Archaerhodopsin in Cholinergic Neurons of Cre-Lox Transgenic Mice. PLoS One 11, e0156596.

Hedrick, T., and Waters, J. (2015). Acetylcholine excites neocortical pyramidal neurons via nicotinic receptors. J Neurophysiol 113, 2195–2209.

Herrero, J.L., Roberts, M.J., Delicato, L.S., Gieselmann, M.A., Dayan, P., and Thiele, A. (2008). Acetylcholine contributes through muscarinic receptors to attentional modulation in V1. Nature 454, 1110–1114.

Joshi, A., Kalappa, B.I., Anderson, C.T., and Tzounopoulos, T. (2016). Cell-Specific Cholinergic Modulation of Excitability of Layer 5B Principal Neurons in Mouse Auditory Cortex. J Neurosci 36, 8487–8499.

Klinkenberg, I., Sambeth, A., and Blokland, A. (2011). Acetylcholine and attention. Behav Brain Res 221, 430–442.

Kolisnyk, B., Al-Onaizi, M.A., Hirata, P.H., Guzman, M.S., Nikolova, S., Barbash, S., Soreq, H., Bartha, R., Prado, M.A., and Prado, V.F. (2013a). Forebrain deletion of the vesicular acetylcholine transporter results in deficits in executive function, metabolic, and RNA splicing abnormalities in the prefrontal cortex. J Neurosci 33, 14908–14920.

Kolisnyk, B., Guzman, M.S., Raulic, S., Fan, J., Magalhaes, A.C., Feng, G., Gros, R., Prado, V.F., and Prado, M.A. (2013b). ChAT-ChR2-EYFP mice have enhanced motor endurance but show deficits in attention and several additional cognitive domains. J Neurosci 33, 10427–10438.

Lange, H.S., Cannon, C.E., Drott, J.T., Kuduk, S.D., and Uslaner, J.M. (2015). The M1 Muscarinic Positive Allosteric Modulator PQCA Improves Performance on Translatable Tests of Memory and Attention in Rhesus Monkeys. J Pharmacol Exp Ther 355, 442–450.

Larkum M. (2013). A cellular mechanism for cortical associations: an organizing principle for the cerebral cortex. Trends Neurosci 36, 141–151.

Magistretti, J., Ma, L., Shalinsky, M.H., Lin, W., Klink, R., and Alonso, A. (2004). Spike patterning by Ca2+-dependent regulation of a muscarinic cation current in entorhinal cortex layer II neurons. J Neurophysiol 92, 1644–1657.

McGaughy, J., Dalley, J.W., Morrison, C.H., Everitt, B.J., and Robbins, T.W. (2002). Selective behavioral and neurochemical effects of cholinergic lesions produced by intrabasalis infusions of 192 IgG-saporin on attentional performance in a five-choice serial reaction time task. J Neurosci 22, 1905–1913.

McGaughy, J., Kaiser, T., and Sarter, M. (1996). Behavioral vigilance following infusions of 192 IgG-saporin into the basal forebrain: selectivity of the behavioral impairment and relation to cortical AChE-positive fiber density. Behav Neurosci 110, 247–265.

Morishima, M., and Kawaguchi, Y (2006). Recurrent connection patterns of corticostriatal pyramidal cells in frontal cortex. J Neurosci 26, 4394–4405.

Nelson, C.L., Sarter, M., and Bruno, J.P. (2005). Prefrontal cortical modulation of acetylcholine release in posterior parietal cortex. Neuroscience 132, 347–359.

Newman, L.A., and McGaughy, J. (2008). Cholinergic deafferentation of prefrontal cortex increases sensitivity to cross-modal distractors during a sustained attention task. The Journal of neuroscience : the official journal of the Society for Neuroscience 28, 2642–2650.

Parikh, V, Kozak, R., Martinez, V, and Sarter, M. (2007). Prefrontal acetylcholine release controls cue detection on multiple timescales. Neuron 56, 141–154.

Parikh V, and Sarter M. (2008). Cholinergic mediation of attention: contributions of phasic and tonic increases in prefrontal cholinergic activity. Ann N Y Acad Sci 1129, 225–235.

Paxinos, G., and Franklin, K.B.J. (2004). The Mouse Brain in Stereotaxic Coordinates, Second edn (San Diego: Academic Press).

Pinto, Y, van der Leij, A.R., Sligte, I.G., Lamme, V.A., and Scholte, H.S. (2013). Bottom-up and top-down attention are independent. J Vis 13, 16.

Porter, J.T., Cauli, B., Tsuzuki, K., Lambolez, B., Rossier, J., and Audinat, E. (1999). Selective excitation of subtypes of neocortical interneurons by nicotinic receptors. The Journal of neuroscience : the official journal of the Society for Neuroscience 19, 5228–5235.

Power, J.M., and Sah, P. (2005). Intracellular calcium store filling by an L-type calcium current in the basolateral amygdala at subthreshold membrane potentials. The Journal of physiology 562, 439–453.

Rahman, J., and Berger, T. (2011). Persistent activity in layer 5 pyramidal neurons following cholinergic activation of mouse primary cortices. The European journal of neuroscience 34, 22–30.

Rasmusson, D.D., Smith, S.A., and Semba, K. (2007). Inactivation of prefrontal cortex abolishes cortical acetylcholine release evoked by sensory or sensory pathway stimulation in the rat. Neuroscience 149, 232–241.

Sarter, M., Lustig, C., Berry, A.S., Gritton, H., Howe, W.M., and Parikh, V (2016). What do phasic cholinergic signals do? Neurobiol Learn Mem 130, 135–141.

Selyanko, A.A., and Brown, D.A. (1996). Regulation of M-type potassium channels in mammalian sympathetic neurons: action of intracellular calcium on single channel currents. Neuropharmacology 35, 933–947.

Seong, H.J., and Carter, A.G. (2012). D1 receptor modulation of action potential firing in a subpopulation of layer 5 pyramidal neurons in the prefrontal cortex. J Neurosci 32, 10516–10521.

Shalinsky, M.H., Magistretti, J., Ma, L., and Alonso, A.A. (2002). Muscarinic activation of a cation current and associated current noise in entorhinal-cortex layer-II neurons. J Neurophysiol 88, 1197–1211.

Ueta, Y, Hirai, Y, Otsuka, T., and Kawaguchi, Y (2013). Direction-and distance-dependent interareal connectivity of pyramidal cell subpopulations in the rat frontal cortex. Frontiers in neural circuits 7, 164.

Ueta, Y., Otsuka, T., Morishima, M., Ushimaru, M., and Kawaguchi, Y. (2014). Multiple layer 5 pyramidal cell subtypes relay cortical feedback from secondary to primary motor areas in rats. Cereb Cortex 24, 2362–2376.

Yan, H.D., Villalobos, C., and Andrade, R. (2009). TRPC Channels Mediate a Muscarinic Receptor-Induced Afterdepolarization in Cerebral Cortex. J Neurosci 29, 10038–10046.

